# Pulmonary Tuberculosis Is Associated with Body Site-Specific Alterations in the Respiratory and Gut Microbiome in Uganda

**DOI:** 10.64898/2026.10.14.759077

**Authors:** Monica Mbabazi, Joanitah Nabwire, Naome Mutesi, Edgar Kigozi, Sharon Namiiro, Adrian Muwonge, David Patrick Kateete

## Abstract

**Background:** Tuberculosis is a major infectious disease whose effects may extend beyond the lungs to influence microbial communities at other body sites. Although studies have reported alterations in the respiratory and gut microbiomes of people with TB, evidence from sub-Saharan Africa remains limited, and the extent to which these alterations differ between body sites is not fully understood. This study characterized sputum and stool microbiomes among Ugandan adults with pulmonary TB and those without TB.

**Methods:** We conducted a cross-sectional study involving 200 participants, comprising 150 adults with pulmonary TB and 50 contacts, at Mulago National Referral Hospital in Kampala, Uganda. Sputum and stool samples were profiled using *16S rRNA* gene sequencing. Following quality control and contaminant removal, within-sample diversity was assessed using multiple alpha diversity indices. Beta diversity was evaluated using distance-based methods, with group differences assessed by permutational multivariate analysis of variance (PERMANOVA) and homogeneity of dispersion tests. Differentially abundant taxa were identified using ANCOM-BC2.

**Results:** After quality control, 135 sputum and 151 stool samples were retained for downstream analysis. In sputum, microbial richness estimators Chao1 and ACE were lower among TB cases than controls, whereas differences in overall community composition were not statistically significant after correction for multiple testing. In stool, several richness and diversity measures differed between groups, and beta-diversity analyses detected differences in community composition. However, TB cases exhibited greater within-group dispersion, and the sequencing platform contributed substantially to variation in stool microbial profiles. In multivariable analyses, the sequencing platform was the only factor independently associated with stool community composition after correction for multiple testing. Differential-abundance analyses identified body site-specific taxonomic associations with TB, including differences involving *Streptococcus*, *Parvimonas*, and *Actinobacillus* in sputum, and *Dorea*, *Lachnoclostridium*, and *Turicibacter* in stool.

**Conclusion:** Pulmonary TB was associated with differences in microbial richness and the abundance of specific taxa in both sputum and stool, but evidence for overall community composition differences varied by body site and was influenced by dispersion and sequencing platform. These findings highlight the importance of examining both respiratory and gut microbial communities in TB research. Longitudinal studies using standardized sequencing methods are needed to clarify how these patterns change during treatment and whether they have diagnostic or clinical relevance.

## Introduction

Tuberculosis (TB), caused by *Mycobacterium tuberculosis* (Mtb), remains a major global public health challenge and the leading cause of death from a single infectious agent. In 2024, an estimated 10.7 million people developed TB and 1.23 million died from the disease worldwide^1^. The burden remains substantial in low- and middle-income countries, including countries in sub-Saharan Africa. Despite advances in diagnosis, treatment, and prevention, important questions remain about the host factors and biological processes that influence TB susceptibility, disease progression, and treatment outcomes. In addition to interactions between Mtb and the host immune system, growing evidence suggests that resident microbial communities may be associated with the pathophysiology of TB and its broader effects on host health^2–5^.

The human respiratory tract harbours diverse microbial communities that contribute to mucosal homeostasis, immune regulation, and resistance to pathogen colonization^6–8^. Disruption of these microbial communities has been implicated in a range of respiratory diseases, including asthma, chronic obstructive pulmonary disease, pneumonia, and tuberculosis^9,10^. Studies investigating the respiratory microbiome of individuals with active TB have reported differences in microbial diversity and shifts in the abundance of specific bacterial taxa^2,5,11,12^. However, findings have been inconsistent across populations and study designs, leaving the extent and consistency of TB-associated respiratory microbiome alterations incompletely understood. Furthermore, focusing exclusively on the respiratory tract may provide an incomplete picture of the microbial changes associated with a systemic infectious disease. Beyond the lungs, emerging evidence suggests that active TB is also associated with differences in gut microbial diversity and composition^13,14^. The gut microbiome plays important roles in host metabolism, immune regulation, and intestinal homeostasis, and alterations in its composition have been linked to several infectious and inflammatory diseases^15–19^. These observations have increased interest in the gut–lung axis, a bidirectional network of interactions through which microbial products, immune signalling, and metabolic pathways may connect the gastrointestinal and respiratory compartments^20,21^. Investigating both sites may therefore provide a broader understanding of the microbial patterns associated with pulmonary TB and their potential relationship to systemic disease processes.

Despite growing interest in the role of the microbiome in tuberculosis, several important knowledge gaps remain. Evidence from sub-Saharan Africa remains limited, and many studies have focused on a single anatomical site, making it difficult to assess whether microbial alterations are observed in the respiratory tract, the gut, or both. Differences in host characteristics, environmental exposures, dietary practices, HIV prevalence, and healthcare access may also influence microbiome composition and limit the generalizability of findings across populations. To address these gaps, we characterized the sputum and stool microbiomes of treatment-naïve Ugandan adults with pulmonary TB and healthy controls using *16S rRNA* gene sequencing. We evaluated microbial richness, diversity, community composition, and differential taxonomic abundance to determine whether active TB is associated with alterations in respiratory and gut microbial communities. By investigating both body sites within a high-TB-burden African setting, this study provides evidence on body site-specific microbiome patterns associated with active TB and contributes to understanding host–microbiome relationships in this population.

## Methods

### Study Design, Site, and Setting

This cross-sectional study investigated differences in the gut and lung microbiota of individuals with pulmonary tuberculosis (TB) and healthy controls in Uganda. Stool and sputum samples were collected and analysed using 16S rRNA gene sequencing to characterize microbial community composition and diversity. Participant recruitment and sample collection were conducted at the Mulago National Referral Hospital TB Clinic in Kampala, Uganda. As the country’s largest national referral and teaching hospital, Mulago serves as a major centre for TB diagnosis, treatment, and research. Laboratory analyses were performed at the Mycobacteriology and Molecular Diagnostics Laboratories, Makerere University College of Health Sciences. These laboratories are CAP-accredited and equipped for molecular microbiology and microbiome research workflows. Bioinformatic and statistical analyses were conducted using high-performance computing resources at the Infectious Diseases Institute, Makerere University, and additional computational infrastructure at the University of Edinburgh.

### Study Population

A total of 200 participants were included in the study, comprising 150 individuals with newly diagnosed pulmonary TB and 50 healthy controls. TB cases were adults aged 18–65 years with pulmonary tuberculosis confirmed using the GeneXpert MTB/RIF assay. Healthy controls were recruited from household contacts and had no clinical evidence of active tuberculosis and confirmed HIV-negative status at the time of enrolment. Following sample processing, sequencing, and quality-control procedures, stool and sputum samples that met predefined inclusion criteria were retained for downstream microbiome analyses. The final number of samples included in each analysis is presented in **Table 1**.

**Table 1.** Summary of sequencing depth across lung (sputum) and gut (stool) samples following QC. Values include total reads, mean reads per sample (± standard deviation [SD]), median reads, and the minimum–maximum range. Sample retention (%) represents the proportion of samples retained after quality filtering relative to the initial number of samples.

| Body Site | Number of Samples (n) | Total Reads | Mean Reads ± SD | Median Reads | Min Reads | Max Reads | Samples retained after QC (%) |
| --- | --- | --- | --- | --- | --- | --- | --- |
| Lung (sputum) | 135 | 1, 054,407 | 7,810 ± 4,694 | 6,597 | 1,399 | 27,678 | 95.1 |
| Gut (stool) | 151 | 292,563 | 2,366 ± 1,007 | 1,969 | 1,204 | 5,896 | 77.4 |

### Sample Collection and Processing

Stool and sputum samples were collected from all enrolled participants at baseline prior to initiation of anti-tuberculosis treatment. Stool samples were used to characterize the gut microbiome, while sputum samples were used as a proxy for the lung microbiome. Sputum specimens were collected using an induced sputum protocol under the supervision of trained clinical personnel. Following sputum induction, participants expectorated respiratory secretions into sterile collection containers for downstream microbiome analyses. Although bronchoalveolar lavage provides a more direct assessment of lung microbiota, it was not used due to its invasive nature. Immediately following collection, samples were transferred into sterile containers containing beta-mercaptoethanol to preserve microbial nucleic acids and minimize degradation during transport. Samples were then transported to the Molecular Biology Laboratory at Makerere University College of Health Sciences and stored at −80°C until DNA extraction. Demographic and clinical information, including age, sex, HIV status, and TB-related clinical characteristics, were collected using structured interviewer-administered questionnaires and standardized clinical data collection forms.

### Sampling Strategy and Sample Size estimation

A convenience sampling approach was used to recruit eligible participants at the Mulago National Referral Hospital TB Clinic. Individuals who met the study eligibility criteria and provided written informed consent were consecutively enrolled until the target sample size was reached. Pulmonary TB cases were identified through routine clinical diagnostic procedures, while eligible household contacts without evidence of active tuberculosis and HIV were recruited as controls. Sample size estimation was performed in R using power-based calculations to ensure adequate statistical power to detect differences in microbial diversity between study groups. For alpha diversity analyses, a two-sample comparison assuming a moderate effect size (Cohen’s *d* = 0.5), a significance level of 0.05, and 90% power indicated a minimum sample size of 85 participants per group. Similarly, power calculations for beta diversity analyses based on PERMANOVA and a moderate effect size (Cohen’s *f* = 0.25) yielded a comparable minimum requirement of approximately 85 participants per group. Based on these calculations, a minimum of 170 participants was required. To increase statistical power, improve the detection of subtle microbiome differences, and accommodate potential sample loss during laboratory processing, sequencing, and quality-control procedures, a total of 200 participants comprising 150 TB cases and 50 healthy controls were enrolled.

### Inclusion and Exclusion Criteria

Participants were eligible for inclusion if they were adults aged 18–65 years with newly diagnosed pulmonary tuberculosis confirmed by GeneXpert MTB/RIF assay, were treatment-naïve at the time of enrollment, and were willing and able to provide written informed consent. For the control group, eligible participants were healthy household contacts of TB patients with no evidence of active TB and confirmed HIV-negative status. Participants were excluded if they had a history of antibiotic use within the two weeks preceding TB diagnosis, as this could significantly alter microbiome composition. Additional exclusion criteria included multidrug-resistant tuberculosis (MDR-TB), severe illness preventing participation in study procedures or provision of informed consent, and pregnancy because of physiological changes that may influence microbial community composition.

### Ethical Statement

This study was conducted in accordance with established ethical standards for research involving human participants. Ethical approval was obtained from the Makerere University School of Biomedical Sciences Research Ethics Committee (SBS-REC), and administrative clearance was granted by Mulago National Referral Hospital. All participants in the parent study had previously provided written informed consent prior to enrollment. Participant confidentiality was strictly maintained through the use of unique study identification numbers, with personal identifiers stored separately from research data. All study procedures adhered to national ethical guidelines and international principles governing research involving human participants.

### DNA Extraction, 16S rRNA Gene Amplification, and Sequencing

Total microbial genomic DNA was extracted from stool and sputum samples using the QIAamp DNA Stool Mini Kit (Qiagen, Hilden, Germany) according to the manufacturer’s instructions. DNA concentration and purity were assessed using a NanoDrop 2000 spectrophotometer (Thermo Fisher Scientific, USA) and a Qubit fluorometer (Invitrogen, USA). Extracted DNA was stored at −80°C until downstream molecular analyses. The V3–V4 hypervariable region of the bacterial 16S rRNA gene was amplified using the universal primers 338F and 806R, generating an amplicon of approximately 460 bp. Polymerase chain reaction (PCR) amplification was performed under standard conditions, after which amplicons were purified and prepared for sequencing using standard library preparation workflows that included adapter ligation, indexing, and magnetic bead-based purification. Library quality and concentration were assessed using a Qubit fluorometer prior to sequencing. Sequencing was performed using paired-end sequencing platforms. Sputum samples and a subset of stool samples were sequenced on the Illumina MiSeq platform using 2 × 300 bp chemistry, while additional stool samples were sequenced on a BGI sequencing platform according to the manufacturer’s protocols.

### Sequence Processing, Taxonomic Assignment, and Data Filtering

Raw sequencing reads were processed using the QIIME 2 bioinformatics platform^22,23^. Sequences were demultiplexed according to sample-specific barcodes and denoised using the DADA2 algorithm, which performs quality filtering, error correction, chimera removal, and inference of amplicon sequence variants (ASVs). Taxonomic classification was performed using a Naïve Bayes classifier trained on the SILVA reference database using a global microbiome dataset generated in our earlier published systematic review and meta-analysis study. Downstream analyses were conducted in R using the *phyloseq* package^24,25^. Metadata were validated and harmonized prior to analysis to ensure consistency between sequencing and clinical datasets. To reduce noise and sparsity, ASVs detected in 10 or fewer samples were excluded prior to downstream analyses. Samples with fewer than 500 sequencing reads following filtering were also removed from the dataset. Non-bacterial sequences, including taxa classified as Eukaryota, Archaea, and unassigned taxa, were excluded. Additional filtering steps were applied to reduce sparsity by removing low-prevalence ASVs and taxa with zero counts following sample filtering. Decontamination procedures were performed prior to downstream analyses.

### Diversity and Community Structure Analyses

Microbial diversity analyses were conducted in R using the phyloseq, vegan, and picante packages. Relative abundance transformation was performed prior to compositional analyses. Alpha diversity was assessed using Observed richness, Chao1 richness, ACE richness, Shannon diversity, Inverse Simpson diversity, Fisher’s alpha, and Faith’s Phylogenetic Diversity (PD). Differences in alpha diversity between groups were evaluated using the Wilcoxon rank-sum test, and p-values were adjusted for multiple testing using the Benjamini–Hochberg (BH) procedure. Adjusted p-values less than 0.05 were considered statistically significant.

Beta diversity was assessed using Bray–Curtis, Jaccard, weighted UniFrac, and unweighted UniFrac distance metrics. Bray–Curtis and weighted UniFrac distances incorporate taxon abundance information, whereas Jaccard and unweighted UniFrac distances are based primarily on taxon presence–absence patterns. Differences in community composition were evaluated using permutational multivariate analysis of variance (PERMANOVA) implemented through the adonis2 function in the vegan package with 999 permutations. Homogeneity of multivariate dispersion was assessed using the betadisper and permutest functions to determine whether observed PERMANOVA results could be attributed to differences in within-group variability. Canonical analysis of principal coordinates (CAP) was used to visualize disease-associated differences in microbial community structure using the capscale function in the vegan package. Unconstrained principal coordinates analyses (PCoA) were additionally performed as supplementary analyses. Because stool samples were generated using both Illumina MiSeq and BGI sequencing platforms and processed independently through denoising and phylogenetic reconstruction workflows, a unified phylogenetic tree was not available for the integrated stool dataset. Consequently, primary beta diversity analyses of stool samples were based on Bray– Curtis and Jaccard distances, while weighted and unweighted UniFrac analyses were conducted separately within each sequencing platform as supplementary analyses.

### Differential Abundance Analysis

Differential abundance analysis was performed at the genus level using ANCOM-BC2^26,27^. Genera exhibiting Benjamini–Hochberg adjusted p-values below 0.05 were considered significantly associated with disease status. Log-fold changes estimated by ANCOM-BC2 were used to identify taxa enriched or depleted among TB cases relative to healthy individuals. Significant taxa were visualized using volcano plots, and abundance patterns were further explored using hierarchical clustering and heatmap analyses.

## Results

### Sequencing output and quality control

Following quality control (QC) and filtering, 1,054,407 sequencing reads were retained from 135 sputum samples, with a mean sequencing depth of 7,810 reads per sample (SD = 4,694; median = 6,597; range = 1,399–27,678). For stool samples, 357,266 sequencing reads were retained across 151 samples, with a reported mean of 2,366 reads per sample (SD = 1,007; median = 1,969; range = 1,204–5,896). The proportions of samples retained after quality control were 95.1% for sputum and 77.4% for stool. Overall, the number of samples retained and sequencing depth differed between the two sample types. Summary statistics for the retained sequencing data are presented in Table 1.

### Alpha Diversity of Lung and Gut Microbiomes

Alpha diversity was assessed to compare within-sample microbial richness and diversity in sputum and stool samples from participants with pulmonary TB (cases) and those without TB (controls) (**Figures 1 and 2**). In sputum samples, TB cases exhibited lower microbial richness than controls, with significantly lower Chao1 and ACE estimates (Figures 1A–B). However, observed richness, Shannon diversity, inverse Simpson diversity, and Fisher’s alpha did not differ significantly between groups. Faith’s phylogenetic diversity was also comparable between cases and controls (Figure 1C). Comparisons stratified by HIV status revealed no statistically significant differences in the richness estimates examined (Figure 1D). Overall, the sputum findings suggest that TB-associated differences in alpha diversity were primarily reflected in richness estimators rather than in broader measures of community diversity.

**Figure 1.**
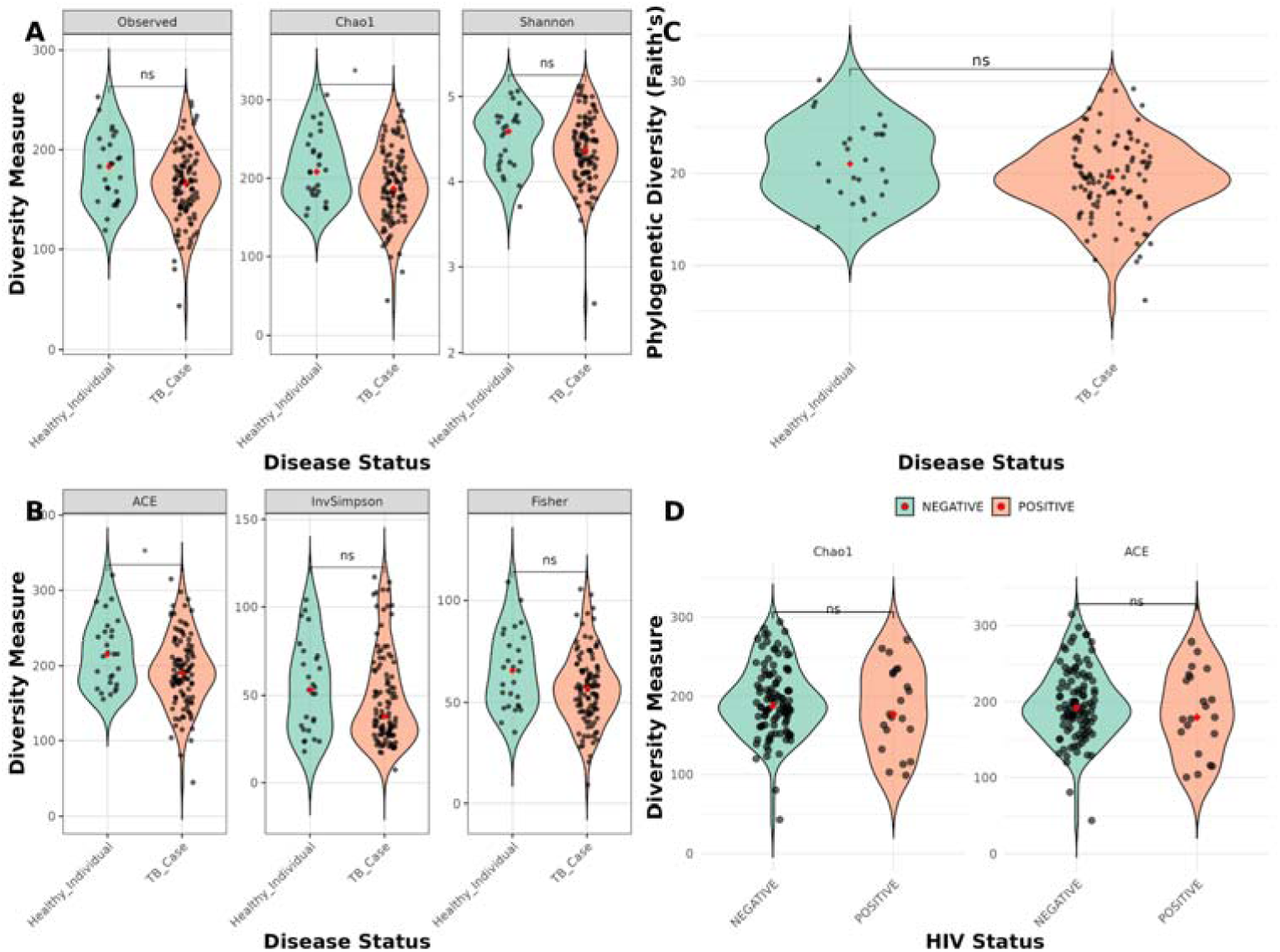
Alpha diversity of the lung (sputum) microbiome stratified by disease and HIV status. (A) Observed richness, Chao1 richness, and Shannon diversity comparing TB cases with healthy controls. (B) ACE richness, inverse Simpson diversity, and Fisher’s alpha comparing the two groups. (C) Faith’s phylogenetic diversity (PD) comparing TB cases with healthy controls. (D) Chao1 and ACE richness estimates stratified by HIV status (HIV-negative vs. HIV-positive). Violin plots show the distributions of diversity measures; overlaid points represent individual samples, and red diamonds indicate group means. Statistical significance is denoted as follows: p < 0.05 (*), p < 0.01 (**), and ns, not statistically significant.

**Figure 2.**
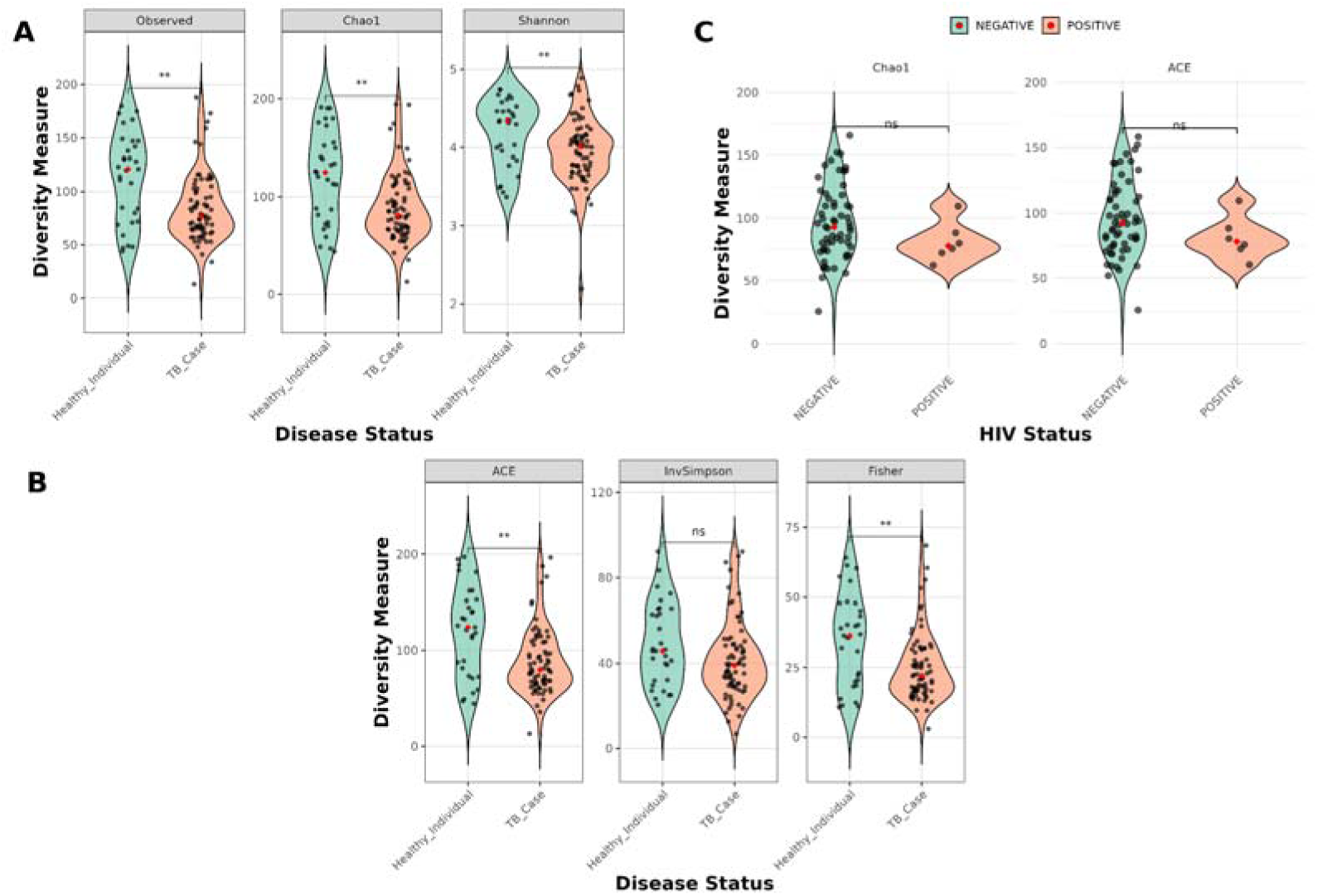
Alpha diversity of the gut (stool) microbiome stratified by TB disease and HIV status. **(A)** Observed richness, Chao1 richness, and Shannon diversity comparing TB cases with healthy controls. **(B)** ACE richness, inverse Simpson diversity, and Fisher’s alpha comparing the two groups. **(C)** Chao1 and ACE richness estimates stratified by HIV status (HIV-negative vs. HIV-positive). Violin plots show the distributions of diversity measures; overlaid points represent individual samples, and red diamonds indicate group means. Statistical comparisons were performed using the Wilcoxon rank-sum test. Statistical significance is denoted as follows: p < 0.05 (*), p < 0.01 (**), and ns, not statistically significant.

In contrast to the relatively modest changes observed in sputum, differences in alpha diversity were more extensive in the gut microbiome (**Figure 2**). TB cases exhibited lower observed richness, Chao1, and ACE estimates than controls (**Figure 2A–B**). Shannon diversity and Fisher’s alpha also differed significantly between groups, whereas the Inverse Simpson index showed no statistically significant difference (**Figure 2B**). Stratification by HIV status revealed no statistically significant differences in the gut microbial richness estimates examined (**Figure 2C**). Overall, these findings suggest that TB-associated differences in gut alpha diversity involved both microbial richness and selected measures of community diversity, whereas differences in sputum were primarily reflected in specific richness estimators. Detailed comparisons of alpha diversity metrics are presented in **Table 2**.

**Table 2.** Alpha diversity metrics of sputum and stool microbiomes stratified by tuberculosis (TB) disease status. Values represent mean alpha diversity estimates for healthy controls and TB cases. Group comparisons were performed using the Wilcoxon rank-sum test. Significance levels are reported as currently available and are subject to confirmation of the Benjamini– Hochberg (BH)-adjusted p-values. Significance is denoted as follows: *p* < 0.05 (*), *p* < 0.01 (**), and ns, not statistically significant.

| Metric | Lung – Controls | Lung – TB cases | p-value | Gut – Controls | Gut – TB cases | p-value |
| --- | --- | --- | --- | --- | --- | --- |
| Observed richness | 180.52 | 163.78 | ns | 132.64 | 131.08 | * |
| Chao1 | 213.29 | 189.94 | * | 141.55 | 137.27 | ** |
| ACE | 216.25 | 192.08 | * | 140.96 | 137.0 | ** |
| Shannon | 4.50 | 4.37 | ns | 4.39 | 4.44 | ** |
| Inverse Simpson | 54.23 | 48.02 | ns | 55.55 | 62.52 | ns |
| Fisher' alpha | 65.58 | 56.96 | ns | 41.92 | 41.36 | ** |
| Faith's PD | 21.30 | 19.49 | ..... | ..... | ..... | ..... |

### Pulmonary TB is associated with modest shifts in sputum microbial community structure

To assess differences in overall sputum microbial community composition between TB cases and controls, beta diversity was evaluated using Bray–Curtis, Jaccard, weighted UniFrac, and unweighted UniFrac distance metrics (Figure 3; Table 3). Canonical analysis of principal coordinates (CAP) showed partial separation between the two groups across all four distance metrics, although substantial overlap remained (Figure 3). The proportion of variation associated with disease status was small, ranging from 1.3% to 2.0%, with the largest value observed for weighted UniFrac (2.0%), followed by Bray–Curtis (1.6%), Jaccard (1.4%), and unweighted UniFrac (1.3%) (Table 3). These findings indicate that disease status accounted for a limited proportion of the variation in sputum microbial community composition, with considerable variation remaining between individuals.

**Figure 3.**
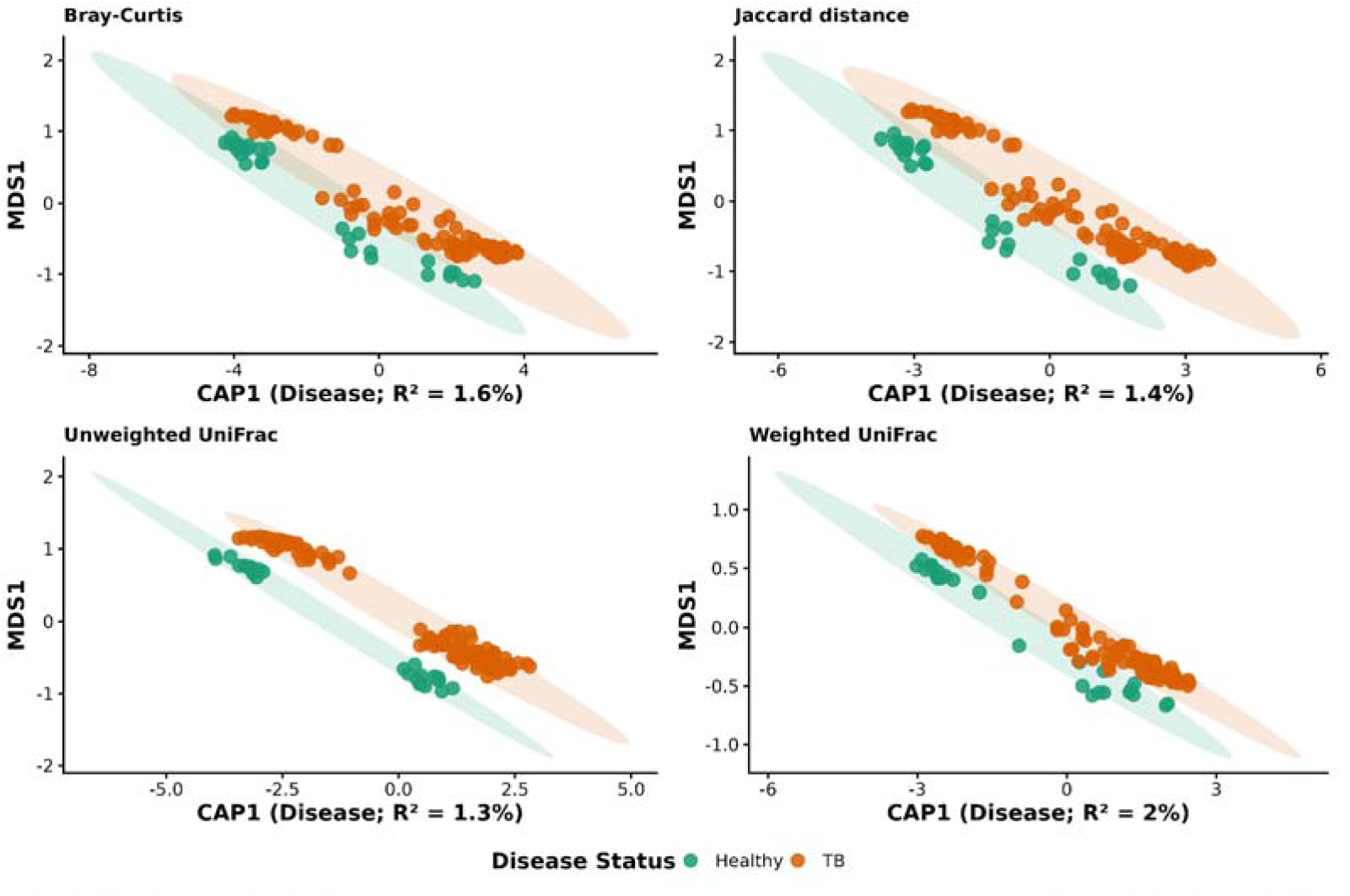
Disease-constrained ordination of sputum microbiome composition in TB cases and healthy individuals. Constrained analysis of principal coordinates (CAP) was performed using Bray–Curtis, Jaccard, unweighted UniFrac, and weighted UniFrac distance metrics to evaluate differences in sputum microbial community composition according to disease status. Each point represents an individual sample, and shaded ellipses indicate the 95% confidence interval for each group. The first constrained axis (CAP1) represents the variation in community composition attributable to disease status, with the proportion of total variation explained (R^2^) indicated on each panel. Disease status explained a modest proportion of the variation in sputum microbiome composition, ranging from 1.3% to 2.0%, with the strongest effect observed for weighted UniFrac distances.

**Table 3.**
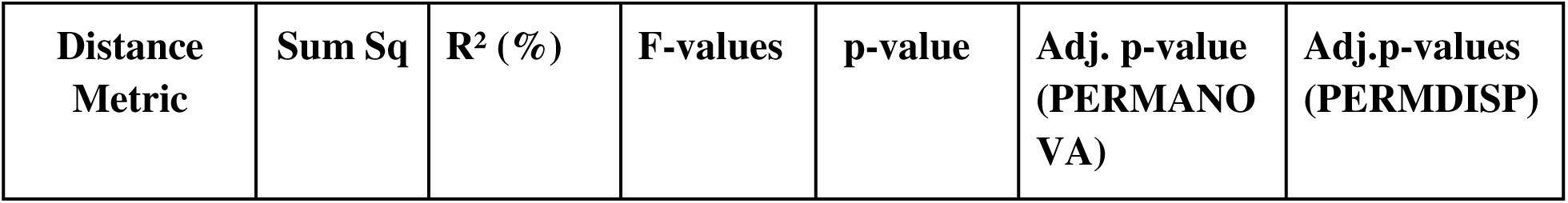
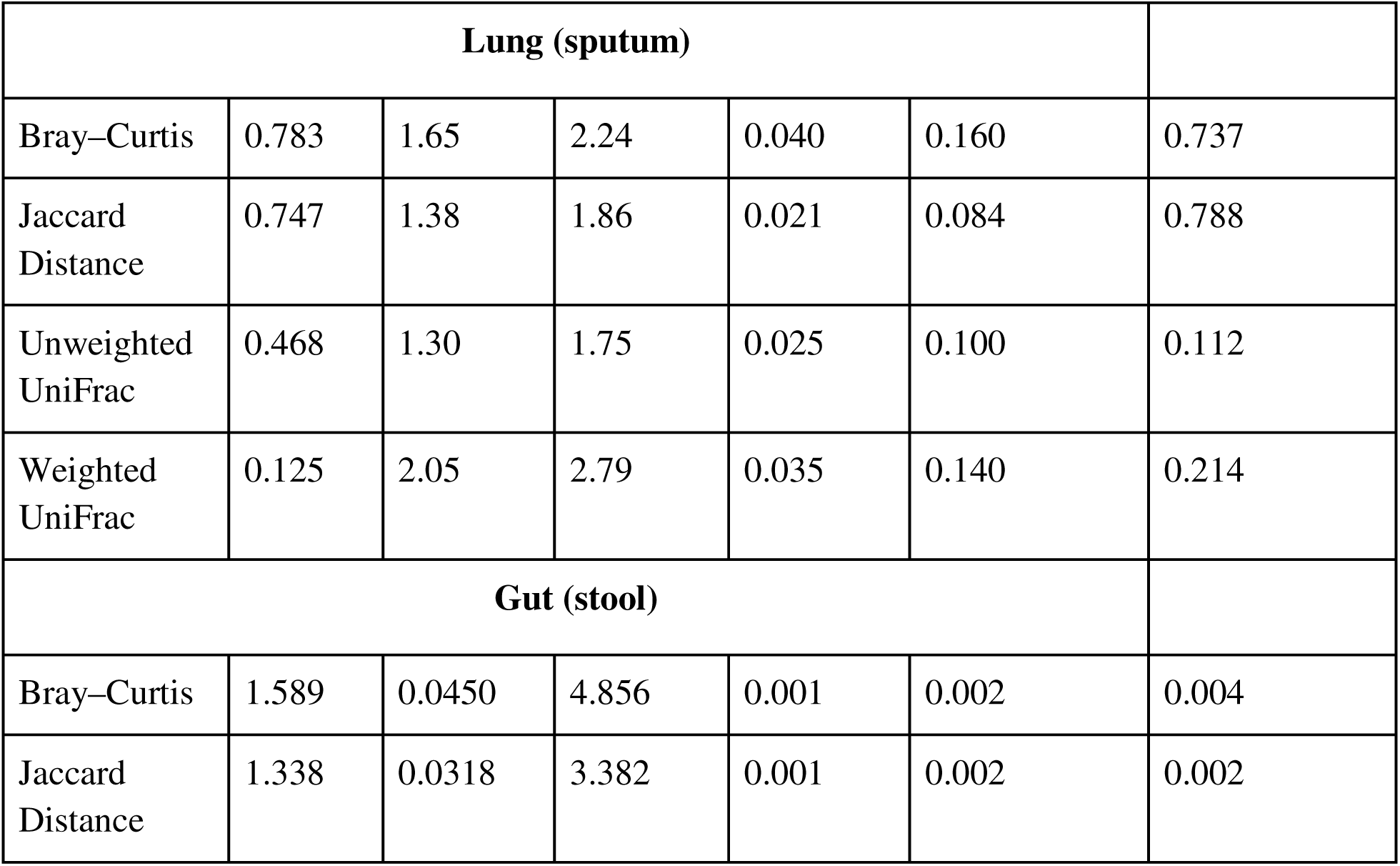
PERMANOVA and PERMDISP analyses across lung and gut microbiomes. Differences in microbial community composition between TB cases and healthy individual were assessed using PERMANOVA based on Bray–Curtis, weighted UniFrac, and unweighted UniFrac distance metrics for both lung (sputum) and gut (stool) microbiomes. Homogeneity of multivariate dispersion was evaluated using PERMDISP. R^2^ values represent the proportion of variation in microbial community structure explained by disease status. Significant PERMANOVA results accompanied by non-significant PERMDISP results were interpreted as robust differences in microbial community composition between groups.

| Distance Metric | Sum Sq | $R^2$ (%) | F-values | p-value | Adj. p-value (PERMANOVA) | Adj.p-values (PERMDISP) |
| --- | --- | --- | --- | --- | --- | --- |

| Lung (sputum) |  |  |  |  |  |  |
| --- | --- | --- | --- | --- | --- | --- |
| Bray–Curtis | 0.783 | 1.65 | 2.24 | 0.040 | 0.160 | 0.737 |
| Jaccard Distance | 0.747 | 1.38 | 1.86 | 0.021 | 0.084 | 0.788 |
| Unweighted UniFrac | 0.468 | 1.30 | 1.75 | 0.025 | 0.100 | 0.112 |
| Weighted UniFrac | 0.125 | 2.05 | 2.79 | 0.035 | 0.140 | 0.214 |
| Gut (stool) |  |  |  |  |  |  |
| Bray–Curtis | 1.589 | 0.0450 | 4.856 | 0.001 | 0.002 | 0.004 |
| Jaccard Distance | 1.338 | 0.0318 | 3.382 | 0.001 | 0.002 | 0.002 |

Unconstrained principal coordinates analysis (PCoA) showed patterns broadly consistent with those observed in the CAP ordination, with substantial overlap between TB cases and healthy controls across the distance metrics examined (Supplementary Figure S3). For Bray–Curtis distances, the first two principal coordinates explained 31.3% and 9.4% of the total variation, respectively. For unweighted UniFrac, the first two coordinates explained 19.4% and 14.9% of the variation, respectively, while the first principal coordinate for weighted UniFrac explained 46.1%. Despite the variation captured by these ordination axes, the plots showed overlapping distributions between disease groups. These findings highlight the substantial inter-individual variability in sputum microbial community composition. To assess whether differences in within-group dispersion could contribute to the observed patterns, homogeneity of multivariate dispersion was evaluated using PERMDISP. No statistically significant differences in dispersion were detected between TB cases and controls for any distance (Supplementary Figure S2). This suggests that unequal within-group dispersion was not detected as an explanation for the observed community patterns.

### Tuberculosis is associated with altered and more heterogeneous gut microbial community structure

Beta diversity was assessed using Bray–Curtis and Jaccard distance metrics to compare gut microbial community composition between TB cases and controls (Figure 4; Table 3). CAP ordination showed partial separation between the two groups for both metrics, although overlap remained (Figure 4). Disease status accounted for 4.4% of the variation in community composition based on Bray–Curtis distances and 3.2% based on Jaccard distances. These findings suggest that TB status was associated with modest differences in gut microbial community composition. PCoA also showed partial separation between TB cases and controls, with variation remaining within both groups. The CAP results indicate that the observed disease-associated differences were modest in magnitude. However, because the Bray–Curtis and Jaccard metrics capture different aspects of community composition, the results should not be interpreted as evidence that changes in the relative abundance of existing taxa predominated over gains or losses of taxa without supporting differential abundance or taxon occurrence analyses.

**Fig 4.**
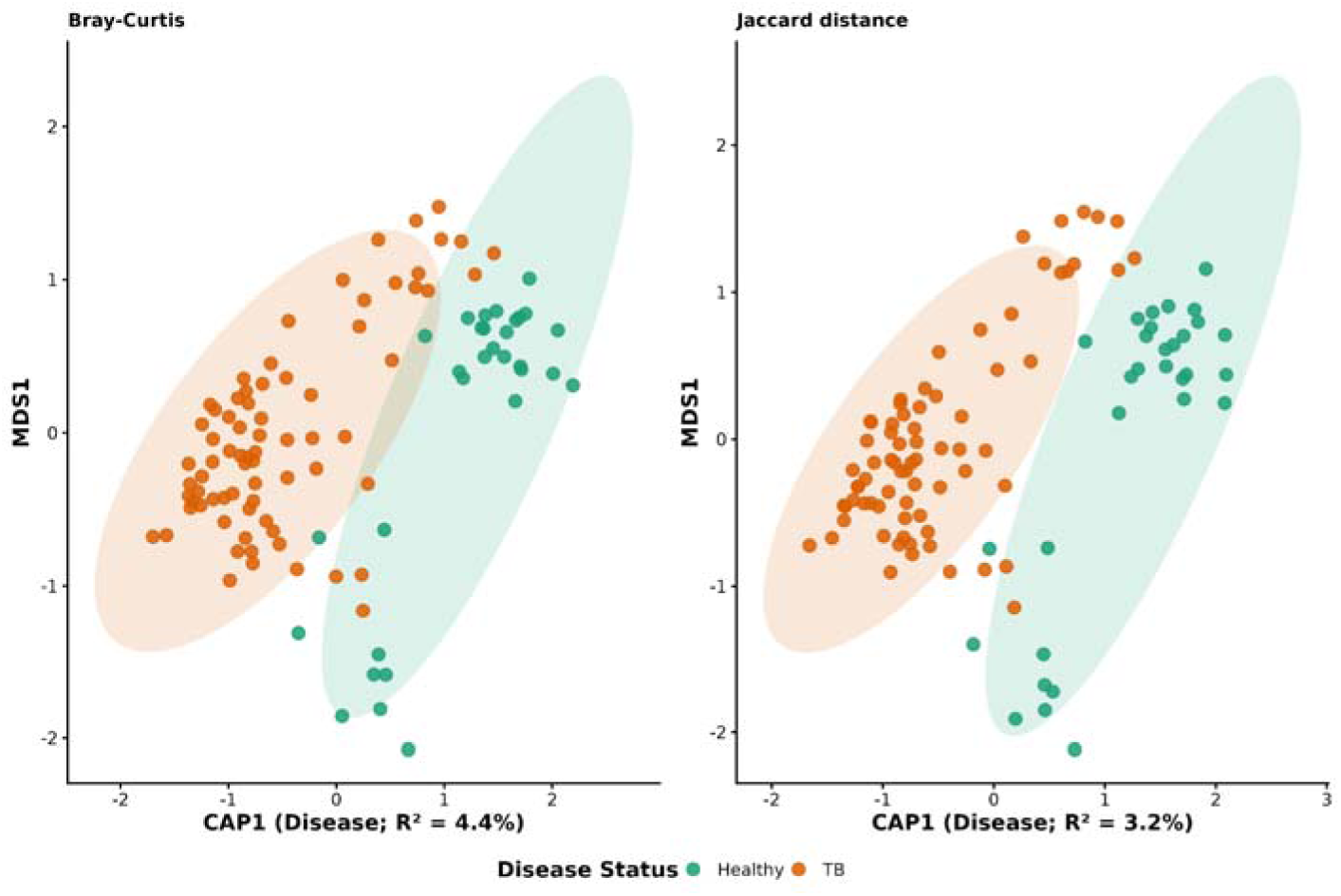
Beta diversity of the gut (stool) microbiome stratified by disease status. Canonical analysis of principal coordinates (CAP) ordinations of gut microbial communities based on **Bray–Curtis** (left) and **Jaccard** (right) distance metrics. Each point represents an individual stool sample and is coloured according to disease status (TB case or healthy individual). Shaded ellipses represent the 95% confidence regions around group centroids. Disease status explained **4.4%** and **3.2%** of the variation in microbial community composition for Bray–Curtis and Jaccard distances, respectively, as indicated by the constrained CAP1 axis. PERMANOVA analyses demonstrated significant differences in overall gut microbial community structure between TB cases and healthy individuals for both distance metrics (Table X). The separation observed along the constrained axis indicates modest but significant disease-associated shifts in gut microbiome composition, with greater dispersion among TB cases suggesting increased inter-individual heterogeneity compared with healthy controls.

To assess whether the observed community patterns were accompanied by differences in within-group variability, homogeneity of multivariate dispersion was evaluated using PERMDISP. TB cases exhibited significantly greater dispersion than healthy controls for both Bray–Curtis and Jaccard distances (adjusted *p* < 0.01 for both comparisons; Supplementary Figures S1 and S2). This finding indicates greater inter-individual heterogeneity in gut microbial community composition among TB cases. Consequently, the PERMANOVA findings should be interpreted alongside the dispersion results, because differences in dispersion can complicate the interpretation of group differences detected by PERMANOVA.

The potential influence of the sequencing platform was also investigated because stool samples had been sequenced using both Illumina MiSeq and BGI platforms before integration for downstream analyses. The sequencing platform was associated with gut microbial community composition, explaining 9.8% and 6.5% of the variation in Bray–Curtis and Jaccard distances, respectively (Supplementary Table S1). In multivariable PERMANOVA, the sequencing platform was the only variable reported to remain independently associated with gut microbial community composition after correction for multiple testing (Supplementary Table S2). Thus, although the unadjusted analyses indicated differences in community composition by TB status, the available multivariable results do not establish disease status as an independent correlate after accounting for the variables included in the model.

Because a unified phylogenetic analysis could not be performed on the combined stool dataset, weighted and unweighted UniFrac analyses were conducted separately within each sequencing platform (Supplementary Figure S4). These analyses showed substantial overlap between TB cases and healthy controls, with limited visual separation between groups. Taken together, the findings indicate that gut microbial community composition differed modestly by TB status in unadjusted analyses, while greater inter-individual heterogeneity among TB cases and the influence of sequencing platform are important considerations in interpreting these differences.

### Taxonomic composition of sputum and stool microbial communities

At the kingdom level, bacterial sequences dominated both sputum and stool microbial communities, accounting for approximately 60% of all classified taxa (**Figure 4A,C**). Eukaryotic taxa represented a smaller proportion of the communities, while a substantial fraction of sequences remained unassigned. Archaeal sequences were rare in both sample types. Overall, the broad kingdom-level composition was highly similar between sputum and stool samples, indicating that bacterial taxa constituted the major component of the microbial communities analysed. Distinct patterns emerged at the phylum level (**Figure 4B,D**). In sputum samples, *Firmicutes* was the most prevalent and abundant phylum, followed by *Bacteroidota, Actinobacteriota, Proteobacteria*, and *Fusobacteriota*. Several additional phyla, including *Patescibacteria, Campylobacterota,* and *Verrucomicrobiota*, were detected at lower prevalence and abundance. In stool samples, Firmicutes was also the dominant phylum, whereas *Actinobacteriota* and *Proteobacteria* represented the next most prevalent taxa. Collectively, these findings indicate that a relatively small number of bacterial phyla accounted for the majority of microbial diversity in both body sites.

Prevalence-abundance analyses revealed a positive relationship between taxon prevalence and average abundance across both sputum and stool microbiomes. Phyla that were detected in a larger proportion of samples generally exhibited higher average abundances, whereas less prevalent phyla tended to occur at lower abundance levels. This pattern was particularly evident for *Firmicutes, Actinobacteriota, Bacteroidota,* and *Proteobacteria*, which were consistently among the most abundant and widely distributed taxa. Together, these results demonstrate that the microbial communities of both sputum and stool samples are structured around a core set of dominant bacterial phyla, accompanied by a larger number of less prevalent and lower-abundance taxa.

### Tuberculosis is associated with shifts in the relative abundance of dominant genera

To investigate taxonomic changes associated with tuberculosis, we compared the relative abundance of the ten most abundant bacterial genera between TB cases and healthy individuals in sputum and stool samples (**Figure 5**). Across both body sites, the dominant genera were largely shared between disease groups, indicating that TB was associated with changes in the abundance of established community members rather than replacement of the core microbiota.

**Figure 5.**
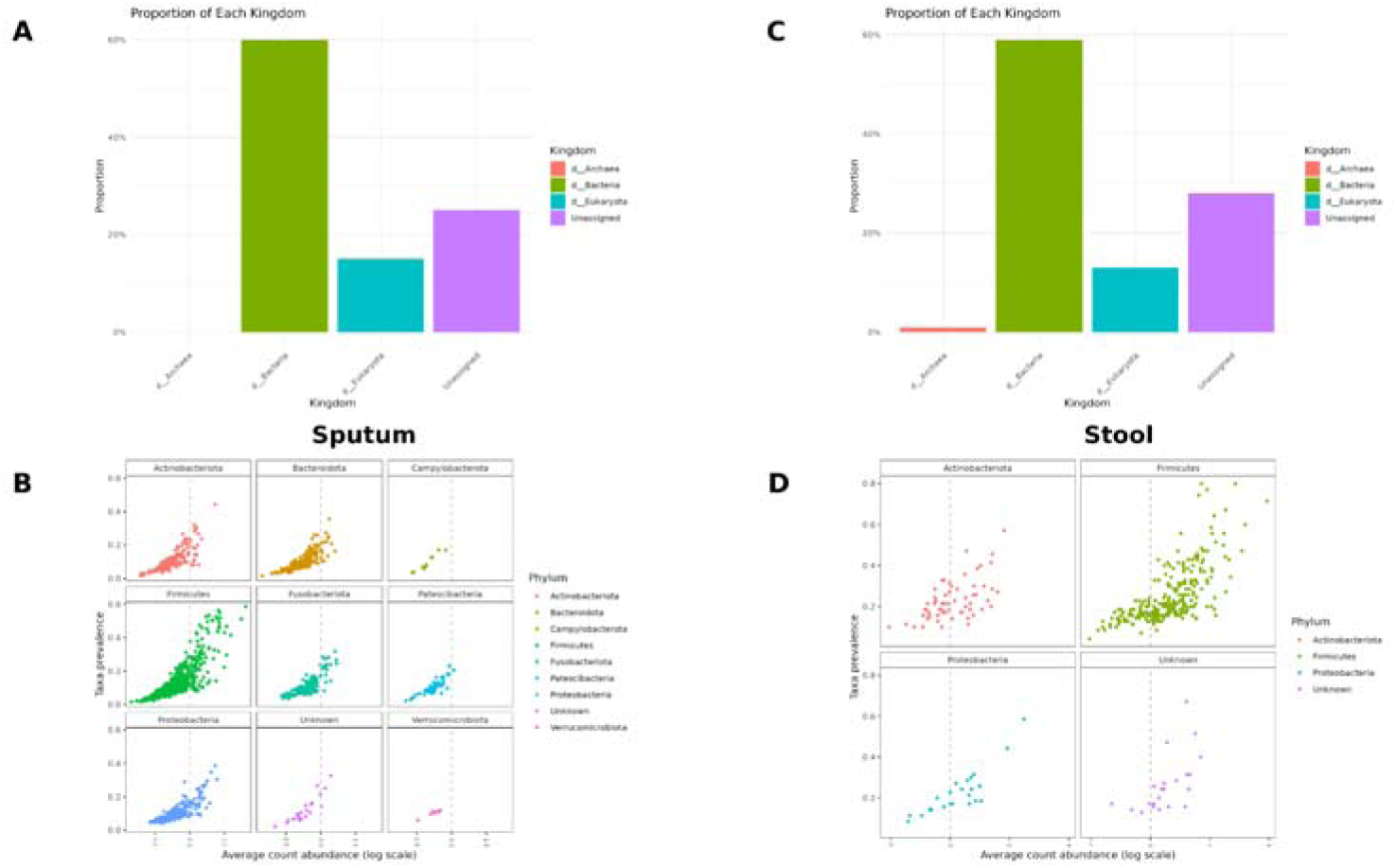
Taxonomic composition and prevalence–abundance distributions of sputum and stool microbial communities. Kingdom-level composition of **(A)** sputum and **(C)** stool microbiomes showed that bacterial sequences dominated both sample types, accounting for approximately 60% of classified taxa, whereas Eukaryota and unassigned taxa represented smaller proportions and Archaea were rare. Phylum-level prevalence–abundance plots for **(B)** sputum and **(D)** stool samples demonstrate a positive relationship between taxon prevalence and abundance. In sputum, Firmicutes, Bacteroidota, Actinobacteriota, Proteobacteria, and Fusobacteriota were among the most prevalent and abundant phyla, whereas stool communities were dominated primarily by Firmicutes, followed by Actinobacteriota and Proteobacteria.

In the sputum microbiome (**Figure 5A**), the community was dominated by *Streptococcus*, *Veillonella*, *Gemella*, *Fusobacterium*, *Leptotrichia*, *Actinomyces*, *Prevotella_7*, *Rothia*, *Neisseria*, and *Conservatibacter*. *Streptococcus* was the most abundant genus in both groups and accounted for a larger proportion of the community among TB cases. In contrast, genera commonly associated with the healthy oral and respiratory microbiota, including *Veillonella*, *Fusobacterium*, and *Neisseria*, appeared relatively reduced in TB cases. Despite these shifts, the overall genus-level composition remained broadly conserved between groups. Similarly, stool microbial communities (**Figure 5B**) were dominated by *Blautia*, *Faecalibacterium*, *Dorea*, *Bifidobacterium*, *Romboutsia*, *Escherichia-Shigella*, and *Subdoligranulum*. While the dominant genera were present in both TB cases and healthy individuals, several compositional differences were evident. TB cases showed relatively higher proportions of *Blautia*, *Dorea*, *Romboutsia*, and *Escherichia-Shigella*, whereas *Faecalibacterium*, *Bifidobacterium*, and *Subdoligranulum* appeared less abundant compared with healthy individuals. These patterns are consistent with the modest but significant differences in gut microbial community structure identified through beta-diversity analyses.

Overall, the genus-level profiles indicate that both lung and gut microbiomes retain a stable core taxonomic composition in the presence of tuberculosis. Disease-associated differences were characterized primarily by shifts in the relative abundance of dominant genera rather than major changes in community membership.

### Distinct body site-specific microbial signatures are associated with tuberculosis

To identify microbial taxa associated with tuberculosis, differential abundance analysis was performed at the genus level using ANCOM-BC2 with Benjamini–Hochberg correction for multiple testing (**Figure 6**). Although beta-diversity analyses indicated only modest shifts in overall community composition, numerous genera were significantly associated with disease status in both stool and sputum samples, suggesting that tuberculosis is characterized by targeted changes in specific microbial taxa rather than wholesale restructuring of microbial communities.

**Fig 6.**
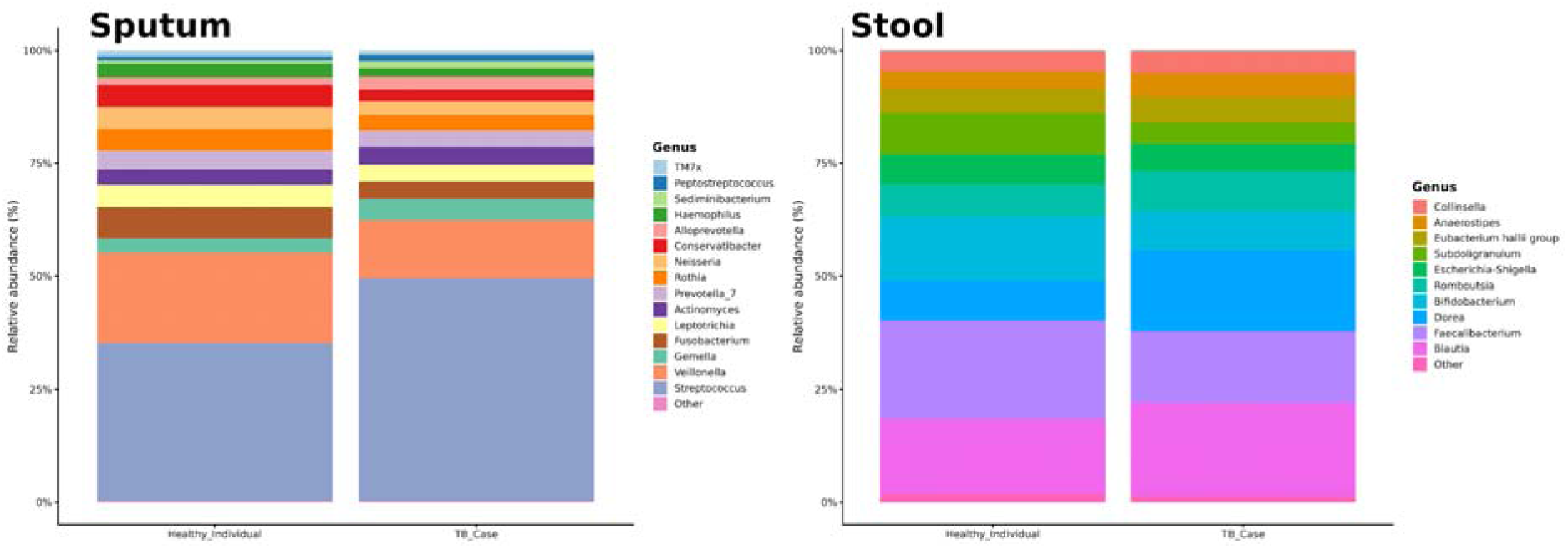
Genus-level taxonomic composition of sputum and stool microbiota in TB cases and healthy individuals. Stacked bar plots showing the mean relative abundance of the ten most abundant bacterial genera identified in sputum (left panel) and stool (right panel) samples. All remaining genera were grouped into the category "Other". Although the dominant genera differed between body sites, the overall genus-level composition within each niche was broadly similar between TB cases and healthy individuals. Differences associated with tuberculosis were primarily reflected by shifts in the relative abundance of dominant taxa rather than the appearance or disappearance of major genera.

Within the stool microbiome, differential abundance analysis identified several genera that differed significantly between TB cases and healthy individuals (**Figure 6, Stool pane**l). Genera enriched among TB cases included *Dorea*, *Lachnoclostridium*, *Intestinibacter*, *Clostridium_sensu_stricto_2*, *Hespellia*, *Erysipelotrichaceae_UCG-003*, *Christensenellaceae_R-7_group*, and *Clostridium_sensu_stricto_1*. Among these, *Dorea* and *Lachnoclostridium* exhibited the largest positive effect sizes and strongest statistical support, indicating a marked increase in abundance among individuals with active tuberculosis. In contrast, *Turicibacter*, *Ruminococcaceae CAG-352*, *Intestinimonas*, *Terrisporobacter*, and *Oscillospiraceae UCG-002* were significantly depleted in TB cases and enriched among healthy individuals. Notably, *Turicibacter* and *Ruminococcaceae CAG-352* showed the strongest negative effect sizes, suggesting that these taxa may represent characteristic features of the healthy gut microbiome that are depleted during active disease. A distinct pattern was observed in the sputum microbiome (**Figure 6, Sputum panel**). Several genera were significantly enriched in TB cases, including *Candidatus_Renichlamydia*, *Actinobacillus*, *Parvimonas*, *Mogibacterium*, *Selenomonadaceae*, *Sphingomonas*, *Lautropia*, *Lachnoanaerobaculum*, *Streptococcus*, and *Alloprevotella*. Many of these taxa are common members of the oral and upper respiratory microbiota and displayed moderate to large positive effect sizes, indicating increased abundance among individuals with active pulmonary tuberculosis. Conversely, several genera were enriched among healthy individuals, including *Corynebacterium*, *Catonella*, *Sediminibacterium*, *Mitochondria*, *Filifactor*, *Neisseriaceae*, and *Uliginosibacterium*. Among these, *Corynebacterium* and *Catonella* exhibited the strongest negative effect sizes and were among the most strongly associated genera in healthy controls.

The differential abundance patterns identified by ANCOM-BC2 were further supported by hierarchical clustering and heatmap analyses. In stool samples, genera enriched in TB cases, including *Dorea*, *Lachnoclostridium*, and *Intestinibacter*, generally exhibited higher relative abundances among TB cases, whereas taxa associated with healthy individuals, such as *Turicibacter* and *Intestinimonas*, were more abundant among controls. Similar patterns were observed in sputum samples, where TB-associated genera such as *Filifactor*, *Atopobium*, *Actinobacillus*, *Gemella*, *Parvimonas*, and *Selenomonadaceae* displayed elevated abundances in subsets of TB samples, while *Uliginosibacterium*, *Sediminibacterium*, *Butyrivibrio*, and *Scardovia* were more frequently associated with healthy individuals.

Collectively, these findings indicate that TB is associated with alterations in both lung and gut microbial communities. While overall community-level differences were modest, particularly in the lung microbiome, tuberculosis was consistently associated with reduced microbial richness, shifts in community composition, and differential abundance of specific taxa. These effects were more pronounced in the gut microbiome, where disease-associated changes were accompanied by increased community heterogeneity.

**Figure 7.**
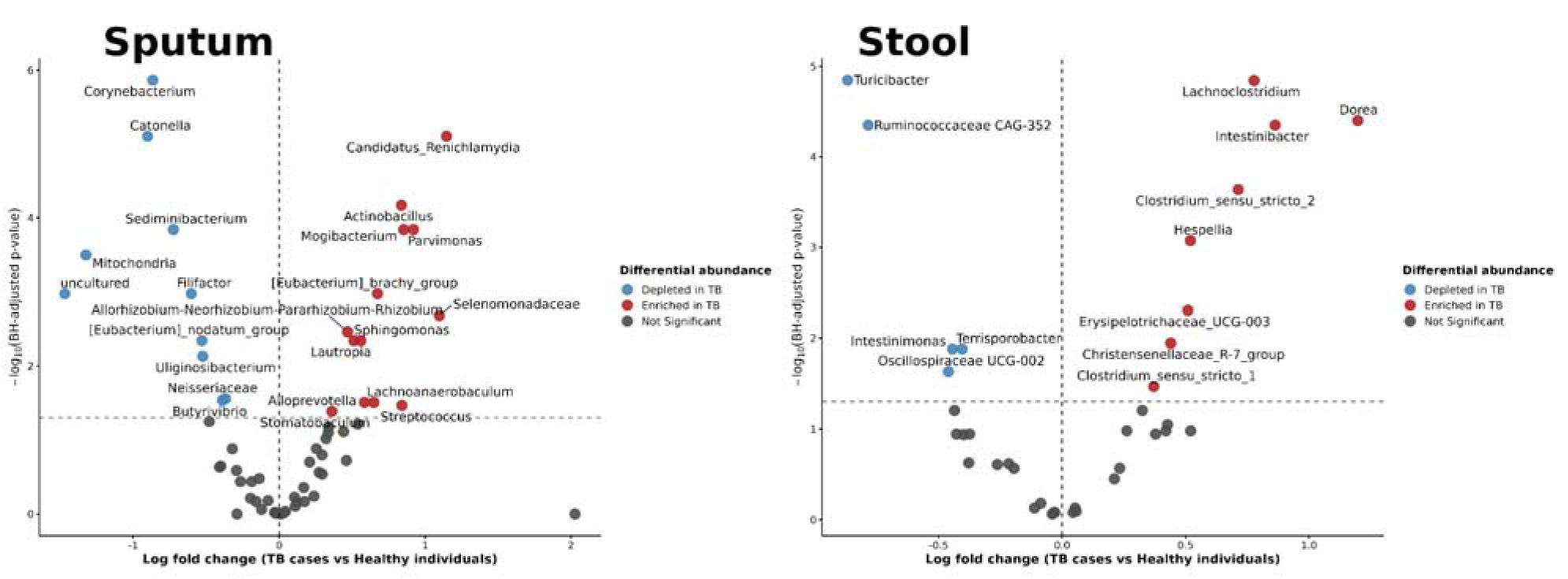
Differentially abundant bacterial genera associated with tuberculosis in sputum and stool microbiomes. Volcano plots showing genus-level differential abundance analysis performed using ANCOM-BC2 with Benjamini–Hochberg correction. The sputum microbiome is shown in the left panel and the stool microbiome in the right panel. Each point represents a bacterial genus, with the x-axis indicating the log fold change between TB cases and healthy individuals and the y-axis representing statistical significance expressed as −log (BH-adjusted p-value). Genera significantly enriched in TB cases are shown in red, genera enriched in healthy individuals are shown in blue, and non-significant genera are shown in grey. Horizontal dashed lines indicate the significance threshold (BH-adjusted p = 0.05), while vertical dashed lines indicate no difference in abundance (log fold change = 0).

## Discussion

### Principal findings

In this study, we characterized the sputum and stool microbiomes of treatment-naïve Ugandan adults with pulmonary TB and controls to investigate disease-associated differences in microbial diversity, community structure, and taxonomic composition. The findings revealed body site-specific patterns of microbial variation associated with active TB. In sputum, TB cases had lower richness estimates, although differences in overall community composition were not statistically significant after correction for multiple testing. In stool, differences in community composition were detected using Bray–Curtis and Jaccard distances; however, these findings require cautious interpretation because TB cases exhibited greater within-group dispersion and the sequencing platform was an important source of variation. Differential abundance analyses also identified taxa associated with disease status in both body sites. Together, these findings suggest that active TB is associated with differences in respiratory and gut microbial profiles, while highlighting the importance of accounting for biological heterogeneity and technical variation when interpreting these associations.

### Body site-specific differences in microbial diversity and community structure

A notable finding of this study was the reduction in microbial richness among individuals with active TB. In sputum samples, richness estimators that account for rare taxa, including Chao1 and ACE, were significantly lower among TB cases, whereas broader measures of diversity such as Shannon and Inverse Simpson indices remained largely unchanged. This pattern suggests that pulmonary TB may be associated primarily with the loss of low-abundance taxa while preserving the overall dominance structure of the respiratory microbial community. Similar observations have been reported in previous respiratory microbiome studies, where active TB was associated with subtle reductions in microbial richness rather than dramatic shifts in overall diversity^5,28,29^. Such changes may reflect the inflammatory environment created by *Mycobacterium tuberculosis* infection, which can alter local ecological conditions within the respiratory tract and reduce the persistence of less abundant microbial populations ^30^.

The impact of TB on the gut microbiome appeared more substantial. In addition to reduced richness, several measures of microbial diversity, including Shannon and Fisher diversity, were significantly lower among TB cases. These findings are consistent with growing evidence that active TB is associated with alterations extending beyond the site of pulmonary infection ^31,32^. The greater magnitude of diversity loss observed in stool samples suggests that systemic effects of TB, including chronic inflammation, altered immune regulation, nutritional changes, and metabolic disturbances, may influence the composition of distal microbial communities ^33^. Reduced gut microbial diversity has been associated with impaired ecosystem stability and diminished resilience to environmental perturbations, raising the possibility that TB-associated dysbiosis may contribute to broader host physiological changes during active disease ^34^.

Differences in microbial community structure were modest and varied by body site. In sputum, disease status explained a small proportion of community variation, and none of the tested beta-diversity comparisons remained statistically significant after correction for multiple testing. This suggests that any TB-associated differences in overall respiratory community structure were relatively subtle in this cohort, despite the taxonomic associations identified through differential abundance analysis. In stool, Bray–Curtis and Jaccard analyses detected differences in community structure between TB cases and healthy controls. However, TB cases also exhibited greater within-group dispersion, indicating greater heterogeneity in gut microbial composition among individuals with active disease. Consequently, the observed gut differences may reflect both shifts in community composition and differences in the variability of microbial communities within groups. In addition, the sequencing platform was an important source of stool microbiome variation, limiting the extent to which the observed differences could be attributed independently to disease status. These findings emphasize the importance of considering effect sizes, group dispersion, and technical variation when interpreting microbiome differences associated with TB.

In contrast, the sputum microbiome exhibited more modest disease-associated differences despite significant alterations in community structure. Several factors may contribute to this observation. First, respiratory microbial communities are inherently dynamic and may be influenced by environmental exposures, oral microbial dispersal, smoking history, and host immune responses ^35^. Second, induced sputum represents a practical but indirect measure of lower respiratory tract microbial communities and may contain contributions from the oral cavity and upper respiratory tract ^36,37^. Although the sputum beta-diversity comparisons did not remain statistically significant after multiple-testing correction, the differential abundance analyses identified specific taxa associated with disease status. These findings suggest that taxonomic differences may be detectable even when overall community-level separation is limited. However, the associations should be interpreted cautiously, particularly given the potential contribution of oral and upper respiratory tract microorganisms to induced sputum samples.

An important observation from our beta-diversity analyses was that disease status explained only a modest proportion of overall microbiome variation, particularly in sputum samples. While these effect sizes were small, they were consistently detected across multiple distance metrics. Microbiomes are influenced by numerous factors, including diet, geography, host genetics, environmental exposures, and socioeconomic conditions, many of which were not directly measured in the present study ^38,39^. Therefore, the relatively low proportion of variation attributable to disease status is expected and likely reflects the multifactorial nature of microbiome in human populations. The consistency of the observed associations across multiple complementary analyses strengthens confidence that the detected microbial differences represent genuine disease-associated signals rather than analytical artefacts.

### Taxonomic differences associated with TB

The community-level patterns observed in the gut microbiome were accompanied by differences in the abundance of specific bacterial taxa. Although the overall taxonomic composition remained broadly similar between TB cases and healthy controls, differential-abundance analysis identified taxa enriched or depleted in association with active disease. These findings suggest that TB-associated gut microbiome alterations may involve shifts in the relative abundance of particular microbial populations rather than wholesale replacement of the gut microbiota. Among the taxa enriched in TB cases, *Dorea* and *Lachnoclostridium* exhibited some of the largest effect sizes and strongest statistical support. Both genera have been associated with altered metabolic and inflammatory states in other human disease contexts ^40,41^. Their enrichment in TB cases may therefore reflect changes in the intestinal microbial environment associated with active disease. However, the precise roles of these taxa in TB remain unclear, and the observed associations do not establish whether their enrichment contributes to disease-related processes or arises as a consequence of changes in host physiology. Previous studies have linked *Lachnoclostridium* and *Dorea* to altered gut microbial profiles in particular disease contexts ^42^, providing a basis for further investigation of their potential relevance to TB-associated microbiome changes. Overall, these findings suggest that active TB is associated with changes in the relative abundance of specific gut microbial taxa, in addition to the differences in diversity observed in this study.

In contrast, several taxa were depleted among TB cases and relatively enriched in controls, including *Turicibacter*, *Intestinimonas*, and members of the families Ruminococcaceae and Oscillospiraceae. These taxa have been associated with gut microbial homeostasis, and some members of these groups have been linked to short-chain fatty acid metabolism ^43^. Their reduced abundance in TB cases may indicate changes in microbial populations associated with intestinal homeostasis, although the functional consequences of these differences were not directly assessed in this study. Together with the taxa enriched among TB cases, these findings suggest that TB-associated gut microbiome differences involve shifts in the relative abundance of specific microbial populations rather than a uniform change across all taxa. However, the cross-sectional design precludes determining whether these microbial differences contribute to altered host immune responses or arise as a consequence of disease-associated physiological changes ^44^.

Distinct taxonomic patterns were also identified within the respiratory microbiome. Although differences in overall community composition between TB cases and individuals without TB were modest and did not remain statistically significant after correction for multiple testing, differential-abundance analyses identified several taxa associated with disease status. These findings suggest that active pulmonary TB may be associated with changes in the relative abundance of specific respiratory microbial populations rather than widespread disruption of the overall community. Taxa enriched among TB cases included *Parvimonas*, *Actinobacillus*, Selenomonadaceae-associated taxa, *Lachnoanaerobaculum*, and *Streptococcus*. Several of these taxa are associated with the oral and upper respiratory tract microbiota ^45^. Their enrichment may reflect changes in the respiratory environment accompanying active disease, although the mechanisms underlying these associations remain uncertain. Conversely, *Corynebacterium*, Neisseriaceae-associated taxa, and *Uliginosibacterium* were relatively enriched among healthy controls. The potential significance of these differences should be interpreted in the context of the ecological roles of the individual taxa and the supporting literature, rather than assuming that their presence necessarily indicates a healthier respiratory ecosystem. Furthermore, induced sputum represents a composite microbial community that may include contributions from the oral cavity and upper respiratory tract. Consequently, the observed taxonomic associations may reflect changes across the broader respiratory niche rather than changes in the lower respiratory tract alone. Taken together, the findings suggest that the gut and respiratory microbiomes exhibit different patterns of variation in association with active TB. Gut community-level differences were more apparent, although these findings require cautious interpretation given the greater within-group dispersion among TB cases and the influence of sequencing platforms. In contrast, the respiratory findings were characterized by associations involving specific taxa despite limited evidence for overall community-level separation. These observations provide a basis for further investigation of body site-specific microbiome patterns and potential systemic influences associated with TB ^46^.

### Methodological considerations

Several methodological considerations are important when interpreting these findings. We evaluated microbial community structure using complementary beta-diversity metrics, which capture different aspects of community variation: Bray–Curtis emphasizes differences in relative abundance, Jaccard focuses on taxon presence or absence, and UniFrac incorporates phylogenetic relationships among taxa. Using multiple metrics provided a broader assessment of community structure, although the results must be interpreted in light of the statistical significance and limitations of each analysis. Canonical analysis of principal coordinates (CAP) was used to visualize variation associated with disease status ^47^, complementing unconstrained principal coordinates analysis (PCoA). However, visual separation in ordination plots should not be interpreted as evidence of statistically significant group differences in the absence of supporting statistical results. Differential abundance was assessed using ANCOM-BC2, which accounts for the compositional nature of microbiome data and incorporates bias correction when estimating taxon-level associations ^48^. Nevertheless, these associations remain observational and require validation in independent cohorts. Finally, examining sputum and stool microbiomes within the same study enabled characterization of microbial patterns across the respiratory and gut compartments, providing a basis for further investigation of potential gut–lung interactions in TB.

### Study limitations

Like any observational microbiome study, our findings should be interpreted in the context of some limitations. First, the cross-sectional design precludes causal inference, making it difficult to determine whether the observed microbial differences contribute to TB susceptibility or arise as a consequence of active disease. Second, respiratory microbial communities were characterized using induced sputum samples, which may contain microbial contributions from the oral cavity and upper respiratory tract and may therefore not fully represent the lower respiratory tract microbiome. Third, stool samples were sequenced using both Illumina MiSeq and BGI platforms. Although platform effects were evaluated in downstream analyses, sequencing platforms remained an important source of variation, and residual technical differences may have influenced the observed microbial profiles. The use of multiple sequencing platforms also precluded construction of a unified phylogenetic framework for the integrated stool dataset, restricting the primary beta-diversity analyses to Bray–Curtis and Jaccard distances. Furthermore, the greater within-group dispersion observed among TB cases in the gut microbiome analyses indicates increased microbial heterogeneity and complicates the interpretation of differences in community composition, which may partly reflect differences in dispersion rather than a uniform shift in microbial composition. Finally, the relatively small number of controls and HIV-positive participants limited statistical power for some subgroup analyses and constrained our ability to investigate interactions between HIV infection, TB, and the microbiome comprehensively. Because participants were recruited from a single TB clinic at Mulago National Referral Hospital in Kampala, the findings may also have limited generalizability to other populations in Uganda and different epidemiological settings.

## Conclusion

This study characterizes body site-specific differences in sputum and stool microbial profiles among Ugandan adults with active pulmonary tuberculosis (TB) and healthy controls. TB cases exhibited differences in microbial richness and the relative abundance of specific taxa, while patterns of overall community composition varied between the two body sites. In sputum, beta-diversity differences did not remain statistically significant after correction for multiple testing. In stool, differences in community composition were detected, although greater within-group dispersion among TB cases and the influence of sequencing platforms warrant cautious interpretation. Together, these findings suggest that TB-associated microbiome patterns extend beyond the respiratory tract and differ between respiratory and gut microbial communities, without establishing whether these differences are causes or consequences of active disease.

These findings contribute to the growing evidence on microbiome alterations associated with TB in a high-burden African setting and highlight the importance of considering both respiratory and gut microbial communities in TB research. Future longitudinal studies using standardized sequencing approaches, larger and more diverse participant groups, and independent validation will be important for clarifying how these microbial patterns change during TB treatment and determining whether the associated taxa have potential diagnostic or clinical relevance.

## Supporting Materials

**Figure S1.**
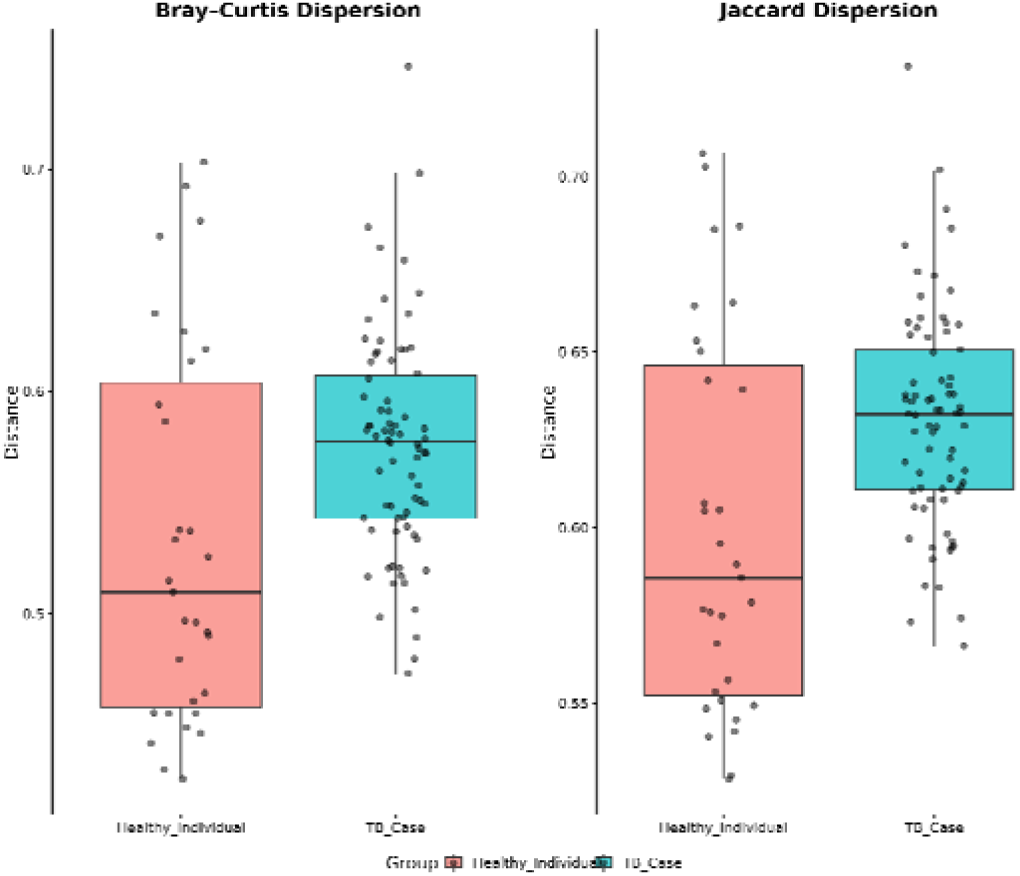
Homogeneity of dispersion (PERMDISP) of gut microbial communities in TB cases and controls. Boxplots show the distances of individual samples to their respective group centroids based on (A) Bray–Curtis and (B) Jaccard dissimilarities. Points represent individual samples. Greater within-group dispersion was observed among TB cases than controls for both distance metrics. The differences in dispersion were statistically significant after multiple-testing correction for Bray–Curtis (adjusted p = 0.004) and Jaccard (adjusted p = 0.002).

**Figure S2.**
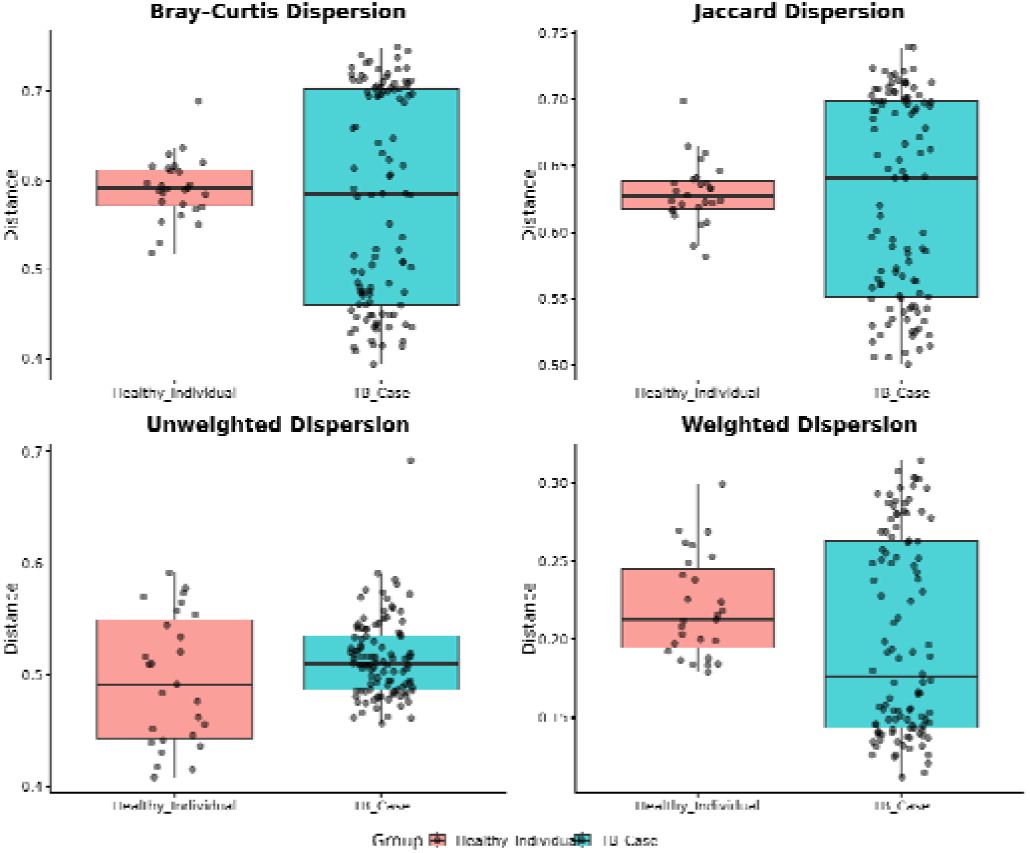
Within-group dispersion of sputum microbial community composition by TB disease status. Homogeneity of multivariate dispersion was assessed using PERMDISP based on Bray–Curtis, Jaccard, unweighted UniFrac, and weighted UniFrac distance metrics. Boxplots show the distribution of distances from individual samples to their respective group centroids for healthy controls and TB cases; overlaid points represent individual samples. The analysis evaluates whether within-group variability in sputum microbial community composition differs by disease status. Statistical significance was assessed using PERMDISP, with p-values adjusted for multiple comparisons where applicable.

**Table 1.**
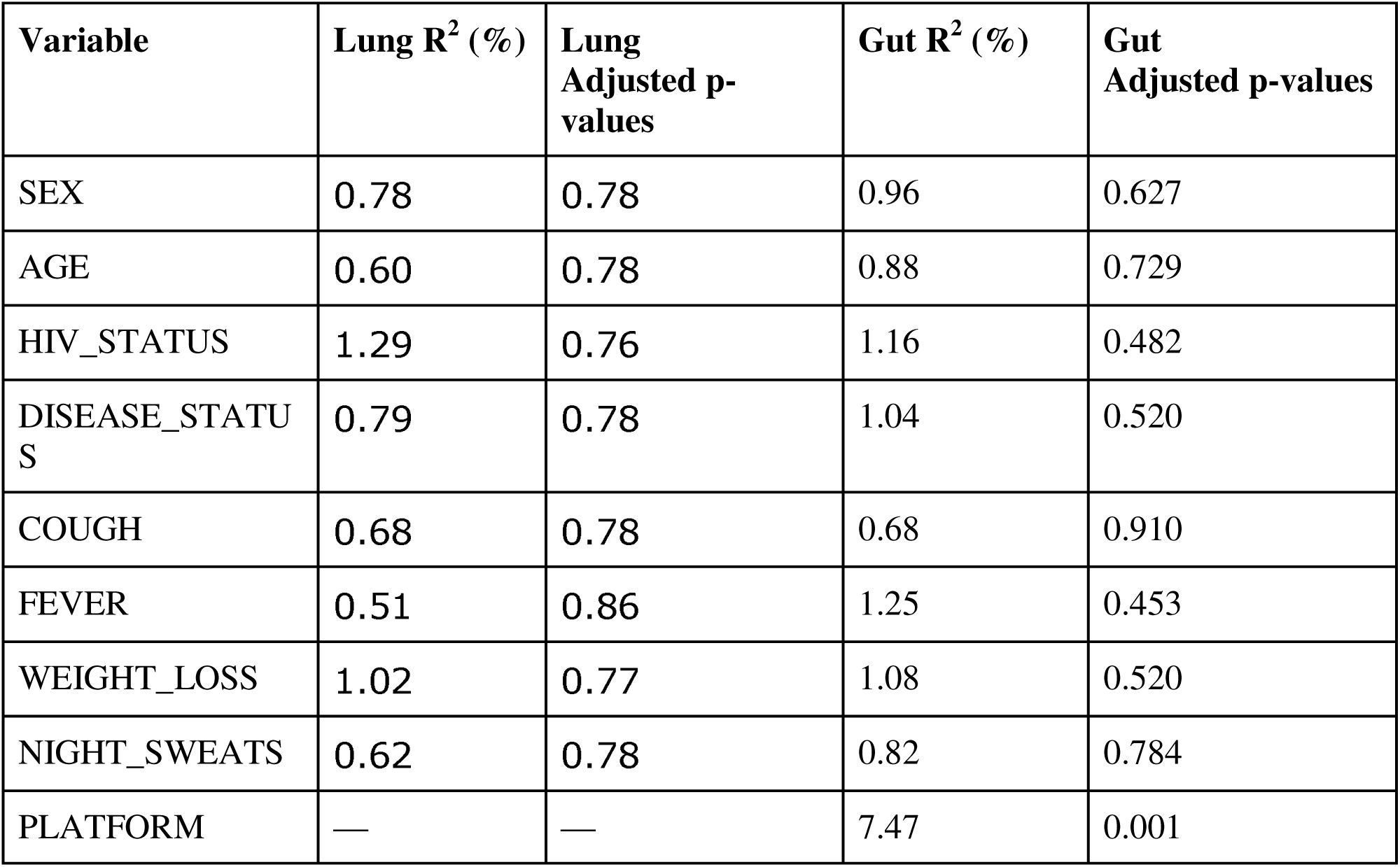
Multivariable PERMANOVA analyses of demographic, clinical, and technical factors associated with lung and gut microbial community composition. Associations between microbial community structure and host characteristics were evaluated using Bray–Curtis distance matrices and multivariable PERMANOVA models. R^2^ values represent the percentage of variation in microbial community composition explained by each variable after accounting for all other variables in the model. P-values were adjusted for multiple comparisons using the Benjamini–Hochberg procedure. For the gut microbiome, sequencing platform (Illumina MiSeq versus BGI) was included as an additional covariate because stool samples were generated using two sequencing platforms. The platform explained the largest proportion of variation in gut microbial community composition (R^2^ = 7.47%, adjusted p = 0.001), whereas no demographic or clinical variable was significantly associated with lung or gut microbial community structure after multiple-testing correction.

**Figure S3.**
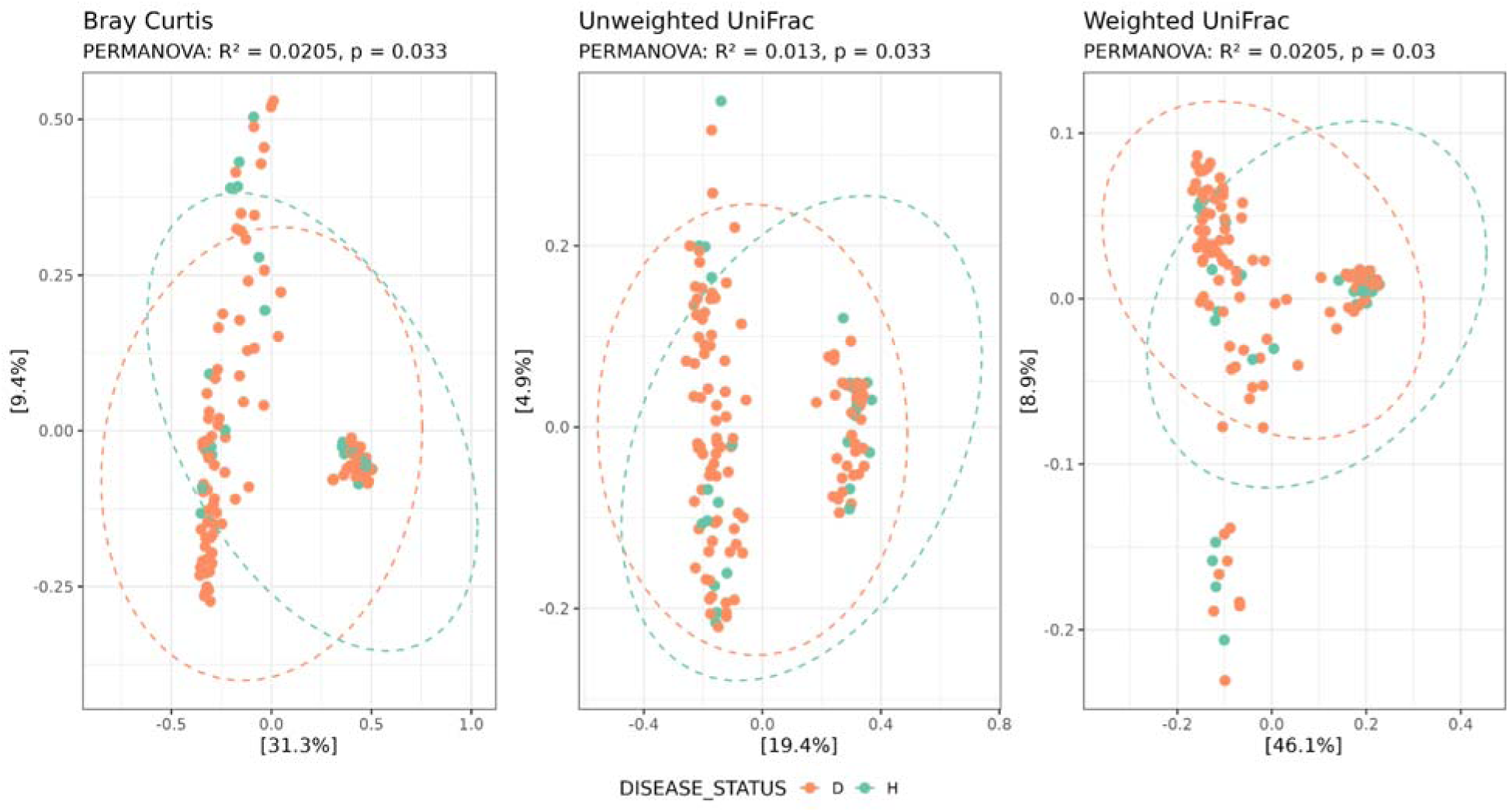
Unconstrained principal coordinate analysis (PCoA) of sputum microbiome composition by tuberculosis (TB) disease status. Non-metric multidimensional scaling (NMDS) ordinations of lung microbial communities based Bray–Curtis, unweighted UniFrac, and weighted UniFrac distance metrics to visualise differences in sputum microbial community composition between TB cases (D, orange) and controls (H, green). Each point represents an individual sample, and dashed ellipses indicate group distributions. The percentages shown on the axes represent the proportion of total variation explained by each principal coordinate. Bray–Curtis, unweighted UniFrac, and weighted UniFrac ordinations showed overlapping distributions between TB cases and healthy controls, indicating substantial inter-individual variation in sputum microbial community composition.

**Figure S4.**
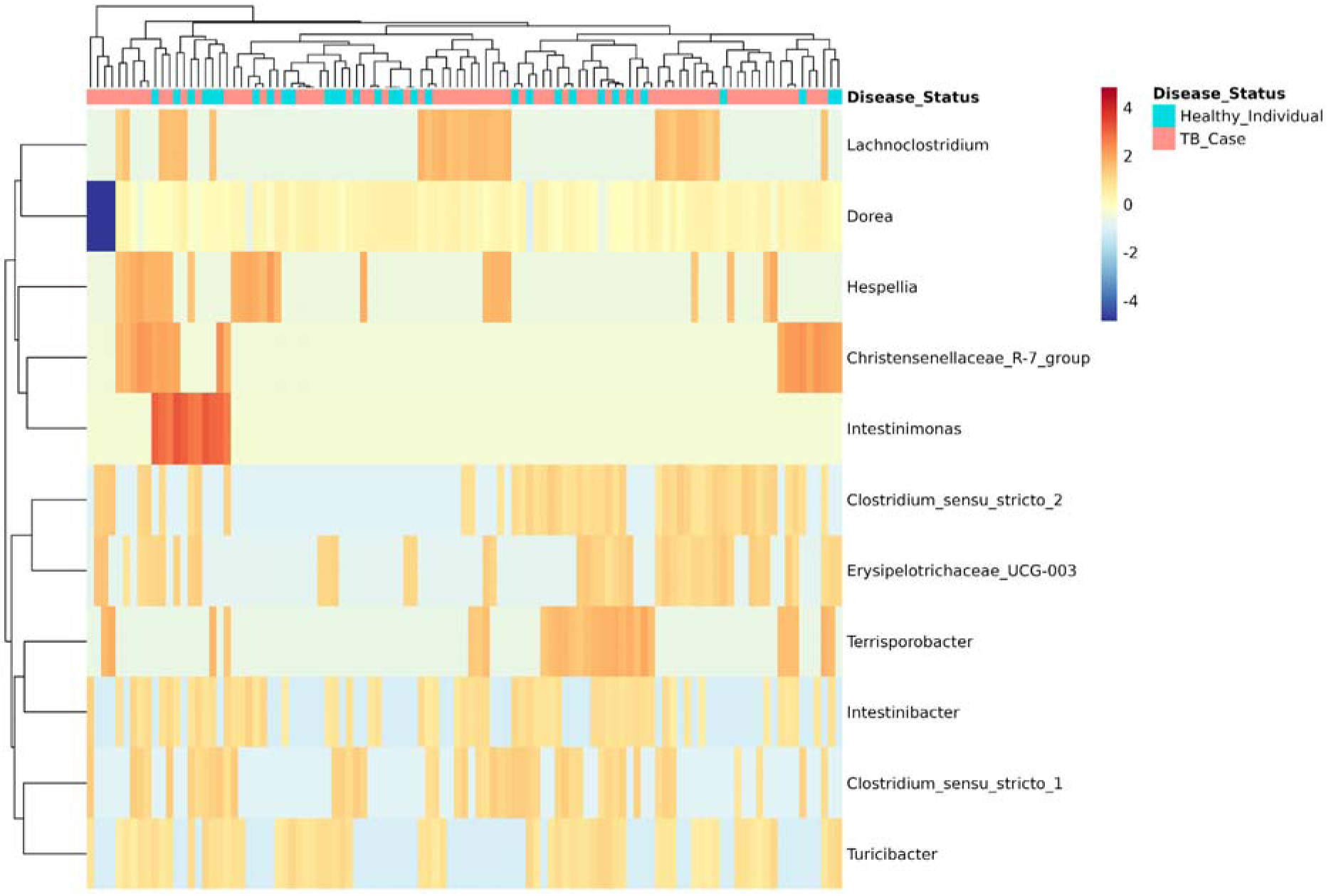
Heatmap of differentially abundant gut microbial taxa identified by ANCOM-BC2. Row-scaled relative abundance profiles of taxa identified as differentially abundant between TB cases and healthy controls are shown across individual stool samples. Columns represent individual samples, and rows represent microbial taxa. The top annotation indicates disease status, and hierarchical clustering was applied to both samples and taxa to visualize similarities in abundance patterns. Red indicates values above the mean for a given taxon, blue indicates values below the taxon-specific mean, and colours near zero indicate values close to the mean. Colour intensity reflects relative variation within each taxon across samples rather than absolute abundance across taxa.

**Figure S5.**
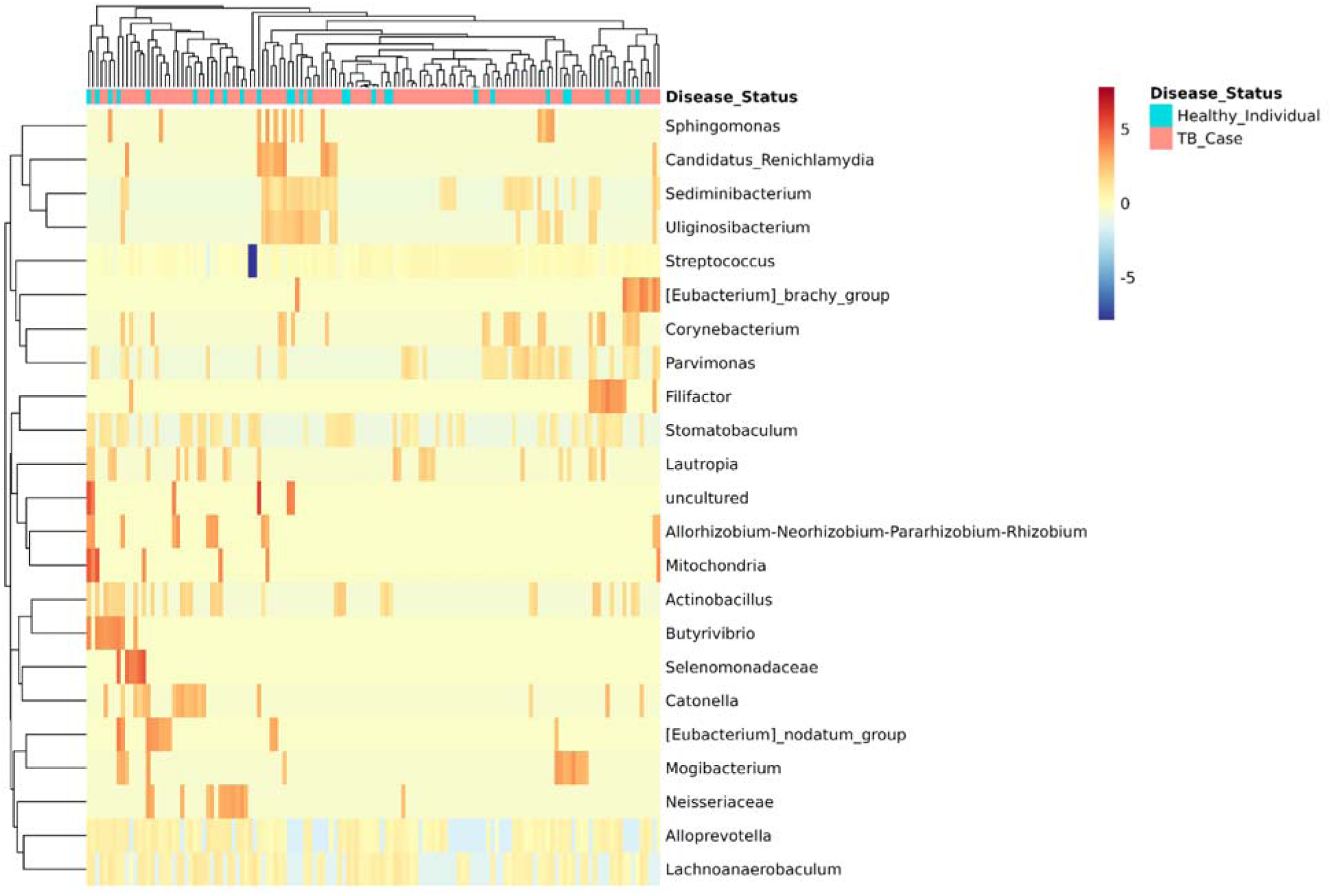
Heatmap of differentially abundant taxa in sputum samples from TB cases and controls. Row-scaled relative abundance profiles of taxa identified as differentially abundant by ANCOM-BC2 are shown across individual sputum samples. Columns represent individual samples, and rows represent microbial taxa. The top annotation indicates disease status, and hierarchical clustering was applied to both samples and taxa to visualize similarities in abundance patterns. Red indicates values above the mean for a given taxon, blue indicates values below the taxon-specific mean, and colours near zero indicate values close to the mean. Colour intensity reflects relative variation within each taxon across samples rather than absolute abundance across taxa. The heatmap illustrates heterogeneous, taxon-specific abundance patterns and substantial inter-individual variation.

