## Supplemental Table 1 for "Pulmonary Tuberculosis Is Associated with Body Site-Specific Alterations in the Respiratory and Gut Microbiome in Uganda"

***Figure S1. Homogeneity of dispersion (PERMDISP) of gut microbial communities in TB cases and controls.***

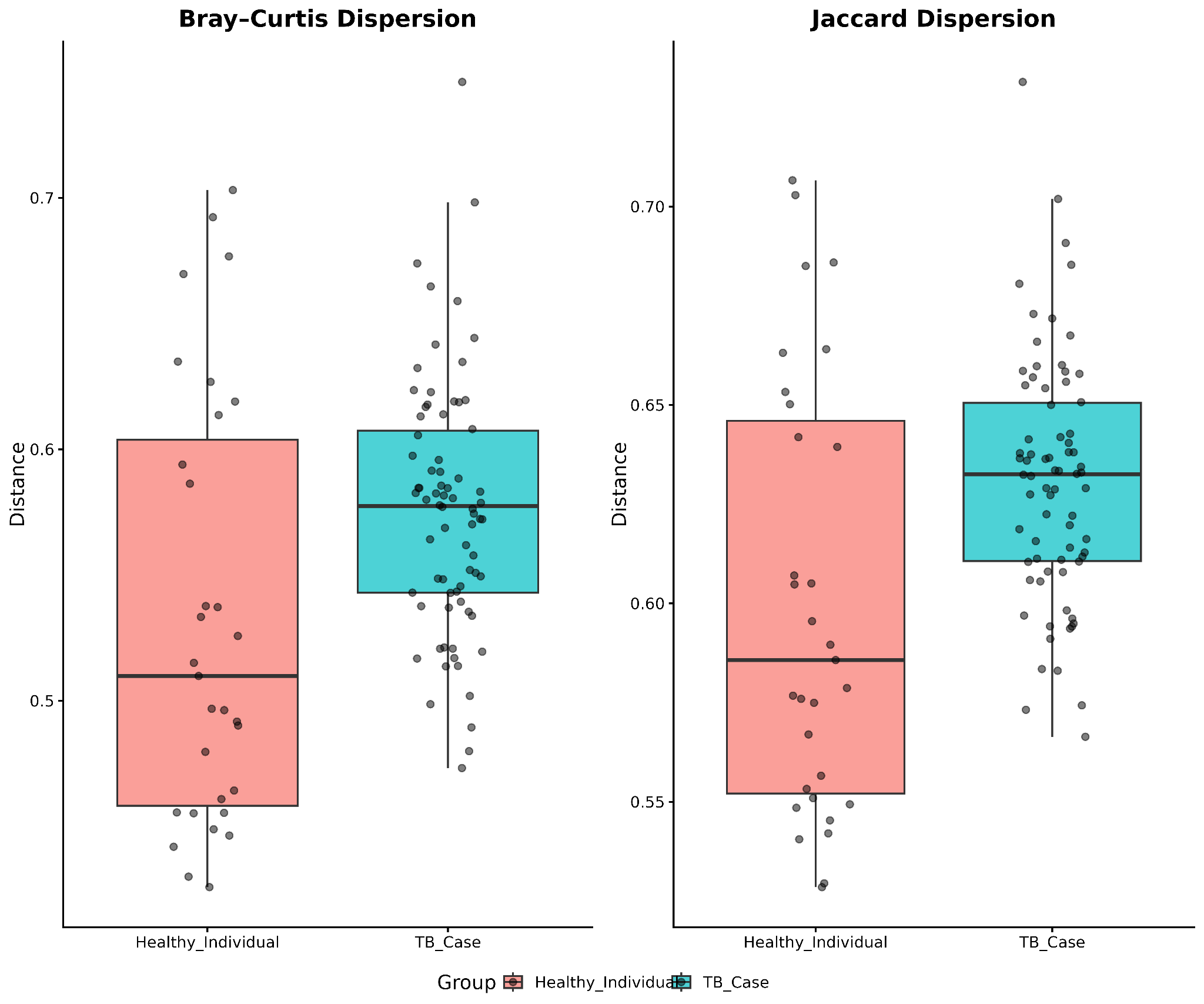

***Figure S1.*** *Boxplots show the distances of individual samples to their respective group centroids based on (A) Bray–Curtis and (B) Jaccard dissimilarities. Points represent individual samples. Greater within-group dispersion was observed among TB cases than controls for both distance metrics. The differences in dispersion were statistically significant after multiple-testing correction for Bray–Curtis (adjusted p = 0.004) and Jaccard (adjusted p = 0.002).*

***Figure S2.*** ***Within-group dispersion of sputum microbial community composition by TB disease status.***

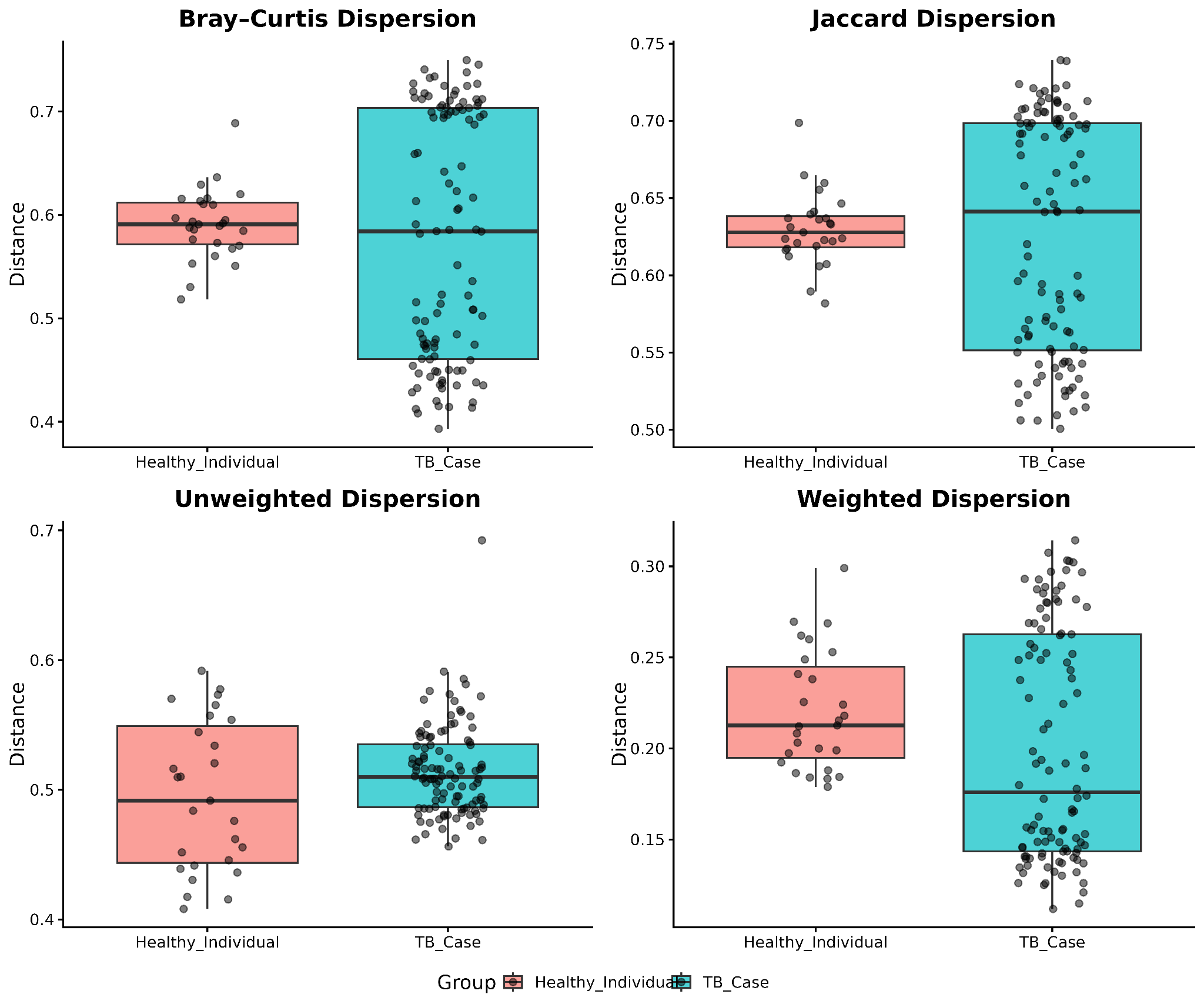

***Figure S2.*** *Homogeneity of multivariate dispersion was assessed using PERMDISP based on Bray–Curtis, Jaccard, unweighted UniFrac, and weighted UniFrac distance metrics. Boxplots show the distribution of distances from individual samples to their respective group centroids for healthy controls and TB cases; overlaid points represent individual samples. The analysis evaluates whether within-group variability in sputum microbial community composition differs by disease status. Statistical significance was assessed using PERMDISP, with p-values adjusted for multiple comparisons where applicable.*

**Table 1. Multivariable PERMANOVA analyses of demographic, clinical, and technical factors associated with lung and gut microbial community composition.**

Associations between microbial community structure and host characteristics were evaluated using Bray–Curtis distance matrices and multivariable PERMANOVA models. R² values represent the percentage of variation in microbial community composition explained by each variable after accounting for all other variables in the model. P-values were adjusted for multiple comparisons using the Benjamini–Hochberg procedure. For the gut microbiome, sequencing platform (Illumina MiSeq versus BGI) was included as an additional covariate because stool samples were generated using two sequencing platforms. The platform explained the largest proportion of variation in gut microbial community composition (R² = 7.47%, adjusted p = 0.001), whereas no demographic or clinical variable was significantly associated with lung or gut microbial community structure after multiple-testing correction.

| **Variable** | **Lung R^2^ (%)** | **Lung**  **Adjusted p-values** | **Gut R^2^ (%)** | **Gut**  **Adjusted p-values** |
| --- | --- | --- | --- | --- |
| SEX | 0.78 | 0.78 | 0.96 | 0.627 |
| AGE | 0.60 | 0.78 | 0.88 | 0.729 |
| HIV_STATUS | 1.29 | 0.76 | 1.16 | 0.482 |
| DISEASE_STATUS | 0.79 | 0.78 | 1.04 | 0.520 |
| COUGH | 0.68 | 0.78 | 0.68 | 0.910 |
| FEVER | 0.51 | 0.86 | 1.25 | 0.453 |
| WEIGHT_LOSS | 1.02 | 0.77 | 1.08 | 0.520 |
| NIGHT_SWEATS | 0.62 | 0.78 | 0.82 | 0.784 |
| PLATFORM | — | — | 7.47 | 0.001 |

***Figure S3. Unconstrained principal coordinate analysis (PCoA) of sputum microbiome composition by tuberculosis (TB) disease status.***

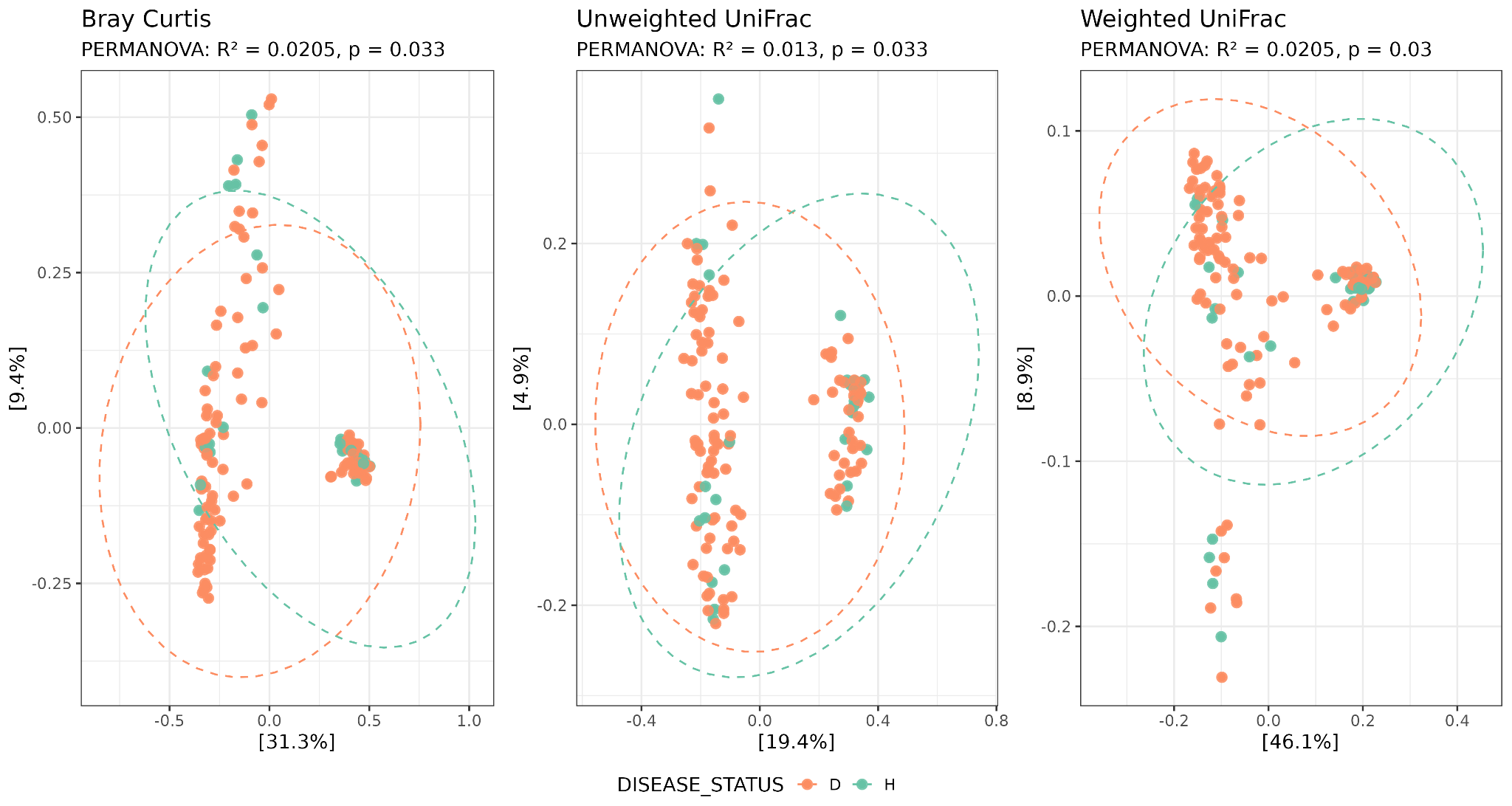

***Figure S3.*** *Non-metric multidimensional scaling (NMDS) ordinations of lung microbial communities based Bray–Curtis, unweighted UniFrac, and weighted UniFrac distance metrics to visualise differences in sputum microbial community composition between TB cases (D, orange) and controls (H, green). Each point represents an individual sample, and dashed ellipses indicate group distributions. The percentages shown on the axes represent the proportion of total variation explained by each principal coordinate. Bray–Curtis, unweighted UniFrac, and weighted UniFrac ordinations showed overlapping distributions between TB cases and healthy controls, indicating substantial inter-individual variation in sputum microbial community composition.*

***Figure S4. Heatmap of differentially abundant gut microbial taxa identified by ANCOM-BC2.***
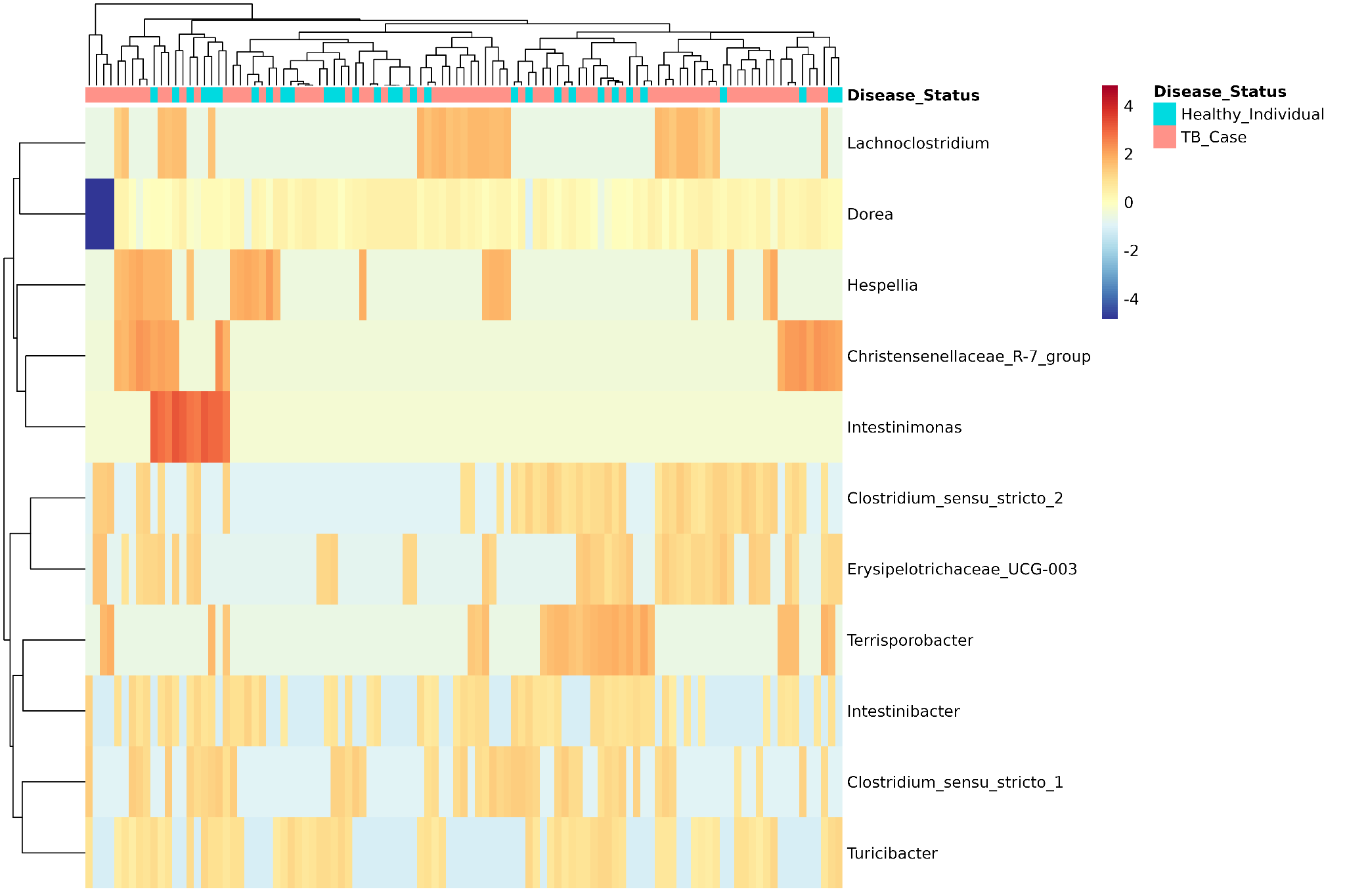

***Figure S4.*** *Row-scaled relative abundance profiles of taxa identified as differentially abundant between TB cases and healthy controls are shown across individual stool samples. Columns represent individual samples, and rows represent microbial taxa. The top annotation indicates disease status, and hierarchical clustering was applied to both samples and taxa to visualize similarities in abundance patterns. Red indicates values above the mean for a given taxon, blue indicates values below the taxon-specific mean, and colours near zero indicate values close to the mean. Colour intensity reflects relative variation within each taxon across samples rather than absolute abundance across taxa.*

***Figure S5. Heatmap of differentially abundant taxa in sputum samples from TB cases and controls.***

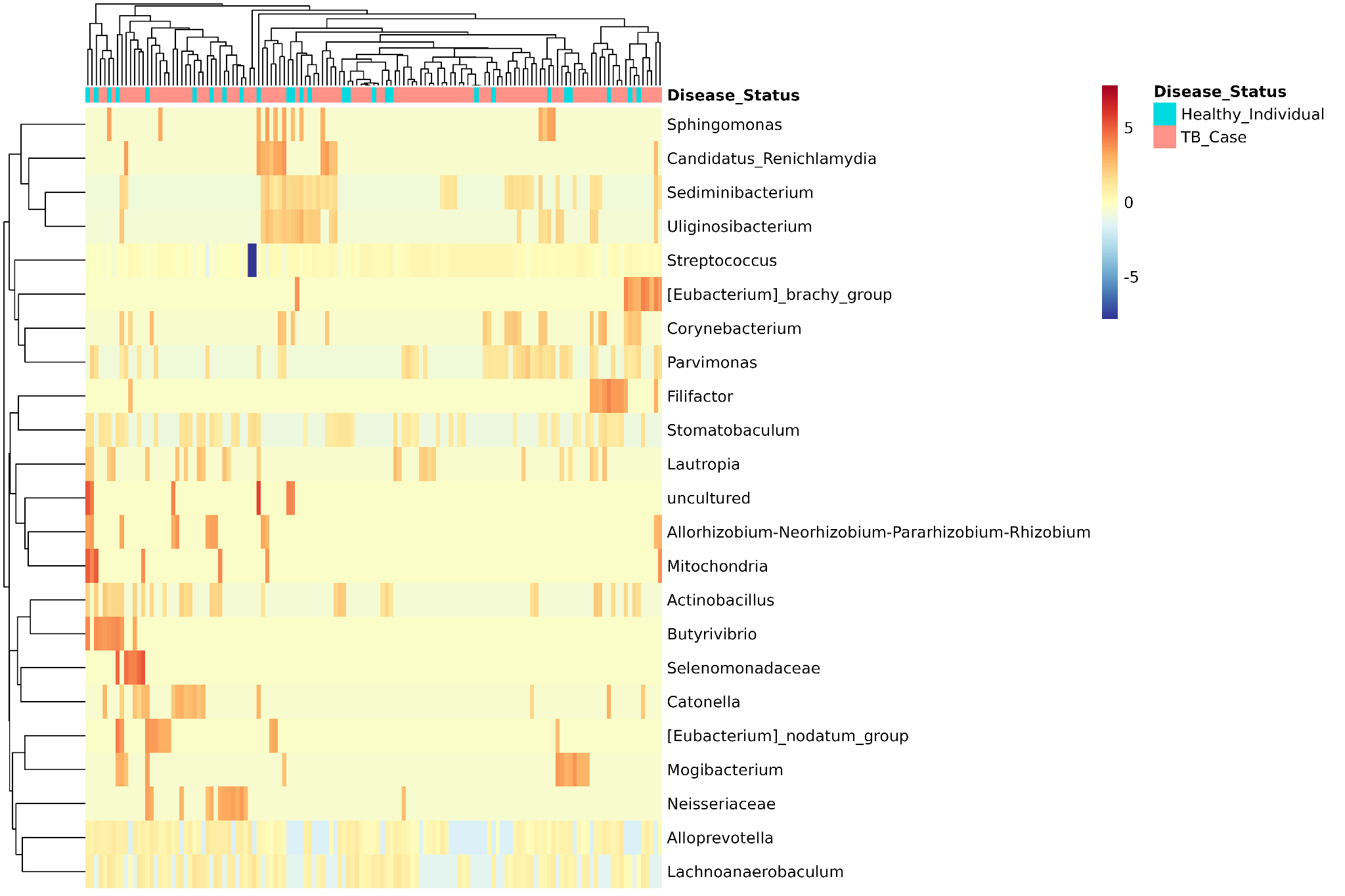
***Figure S5.*** *Row-scaled relative abundance profiles of taxa identified as differentially abundant by ANCOM-BC2 are shown across individual sputum samples. Columns represent individual samples, and rows represent microbial taxa. The top annotation indicates disease status, and hierarchical clustering was applied to both samples and taxa to visualize similarities in abundance patterns. Red indicates values above the mean for a given taxon, blue indicates values below the taxon-specific mean, and colours near zero indicate values close to the mean. Colour intensity reflects relative variation within each taxon across samples rather than absolute abundance across taxa. The heatmap illustrates heterogeneous, taxon-specific abundance patterns and substantial inter-individual variation.*
